# Microtubule competition and cell growth recenter the nucleus after anaphase in fission yeast

**DOI:** 10.1101/2023.01.31.526443

**Authors:** Kimberly Bellingham-Johnstun, Annelise Thorn, Julio Belmonte, Caroline Laplante

## Abstract

Cells actively position their nucleus based on their activity. In fission yeast, microtubule-dependent nuclear centering is critical for symmetrical cell division. After spindle disassembly at the end of anaphase, the nucleus recenters over a ~90 min period, approximately half of the duration of the cell cycle. Live cell and simulation experiments support the cooperation of two distinct mechanisms in the slow recentering of the nucleus. First, a push-push mechanism acts from spindle disassembly to septation and involves the opposing actions of the mitotic Spindle Pole Body microtubules that push the nucleus away from the ends of the cell while post-anaphase array of microtubules basket the nucleus and limit its migration toward the division plane. Second, a slow-and-grow mechanism finalizes nuclear centering in the newborn cell. In this mechanism, microtubule competition stalls the nucleus while asymmetric cell growth slowly centers it. Our work underlines how intrinsic properties of microtubules differently impact nuclear positioning according to microtubule network organization and cell size.

## INTRODUCTION

Active nuclear positioning through microtubule dynamics is an essential process for many types of cells (Gundersen and Worman, 2013). In fission yeast, the nucleus localizes to the cell center during interphase and instructs the position of the contractile ring to ensure symmetrical division (Tran *et al*., 2000; Tran *et al*., 2001; Daga *et al*., 2006). A microtubule-dependent mechanism centers the nucleus during interphase ahead of cytokinesis. Bundles of interphase microtubules attached to the nuclear envelope at the Spindle Pole Body (SPB) and other sites polymerize until they reach the cell ends (Ding *et al*., 1997; Hagan and Yanagida, 1997; Tran *et al*., 2001). Their polymerization against the end of the cell generates pushing forces that are transmitted to the nucleus and push it away from that cell end. Cycles of microtubule polymerization followed by catastrophe pushing at both cell ends result in the oscillation of the nucleus and in its directed recentering when the nucleus is artificially moved off center (Tran *et al*., 2001; Daga *et al*., 2006). Simulations of this nuclear positioning mechanism constrained by microtubule dynamics measured in live cells were sufficient to recapitulate the oscillatory and directed motions of the nucleus observed in late interphase cells (Tran *et al*., 2001; Daga *et al*., 2006; Foethke *et al*., 2009).

During anaphase, the daughter nuclei are naturally displaced away from the cell center and pushed to the cells ends by the spindle. How and when the nuclei reposition back to the cell center after spindle disassembly remains unknown. The Post-Anaphase Array of (PAA) microtubules and the mitotic Spindle Pole Body (mSPB) microtubules are potential candidates for repositioning the nucleus after the end of anaphase. The PAA microtubules form a ring-like structure associated with the contractile ring (Hagan and Hyams, 1988; Pichova *et al*., 1995; Heitz *et al*., 2001; Pardo and Nurse, 2003). They polymerize from equatorial MicroTubule Organizing Centers (eMTOCs) recruited to the contractile ring in myosin-II Myp2-dependent manner (Samejima *et al*., 2010). Different roles have been proposed for these microtubules including anchoring the contractile ring, reestablishing interphase microtubules before the end of cytokinesis and keeping the nucleus away from the ingressing septum (Hagan and Yanagida, 1997; Pardo and Nurse, 2003). The mSPB microtubules assemble from the SPB after spindle disassembly (Samejima *et al*., 2010). Their specific role has not been experimentally determined but interpretation of fixed and immunostained cells have suggested that the mSPB microtubules coordinate with the PAA microtubules to pull the nucleus back to the cell center after anaphase (Hagan and Yanagida, 1997).

We found that the nucleus is off center from spindle disassembly until septation, a period lasting ~50 min, and eventually recenters ~40 min after septation. We identified two microtubule-based mechanisms that reposition the nucleus during this ~90 min period. First, a push-push mechanism involving two competing networks of microtubules position the nucleus at a third of the distance between the cell end and the contractile ring before septation. The mSPB microtubules push the nucleus away from the cell end toward the constricting contractile ring while the PAA microtubules push the nucleus in the opposite direction. After septation, cell growth in combination with microtubules slowly complete the centering of the nucleus. Simulations constrained by microtubule dynamics show that microtubule competition in the short newborn cell is responsible for the slow velocity of nuclear displacement. Importantly, the addition of asymmetric cell growth to the simulation was necessary to explain the profile of nuclear displacement after septation supporting a slow-and-grow mechanism of nuclear displacement in the newborn cell. Our work establishes that the same intrinsic properties of microtubules in cells of different sizes, and perhaps shape, can result in different outcomes on the positioning of the nucleus.

## RESULTS

### Distinct microtubule networks recenter the nucleus after anaphase

At the onset of mitosis, the spindle elongates between the duplicated SPBs and drives the daughter nuclei, hereafter referred to as nuclei, to the opposite ends of the cell (Supplemental Figure 1A). To determine when the nucleus recentered, we tracked the position of the nucleus from the time of spindle disassembly in time-lapse micrographs of cells expressing either Cut11p-tdTomato (nuclear marker), GFP-Atb2p (microtubules) and Sad1p-mEGFP (SPB), or Sad1-RFP and mEGFP-Myo2p (contractile ring marker). Coincident with spindle disassembly, the nucleus started to reposition toward the center of the half-cell compartment (Figure 1A). The nucleus moved in a directed manner until it reached its destination and then oscillated about that position. In wild-type cells, the destination of the nucleus, the maximal distance of the center of the nucleus from the cell end, was 2.3 ± 0.4 μm (mean ± standard deviation) and from the contractile ring was 3.7 ± 0.6 μm (n = 20 nuclei) (Figure 1A and B). This position corresponds to a third of the distance between the end of the cell to the contractile ring. After septation, in the newborn cell, the nucleus resumed its motion after ~10 min and reached a stable position just beyond the cell center ~40 min after septation (Figure 1B and C). The total displacement of the nucleus thus happened in two phases with the first phase spanning the end of anaphase until septation and the second phase occurring over the first 40 min in the newborn cell for a total duration of ~90 min.

**Figure 1.**
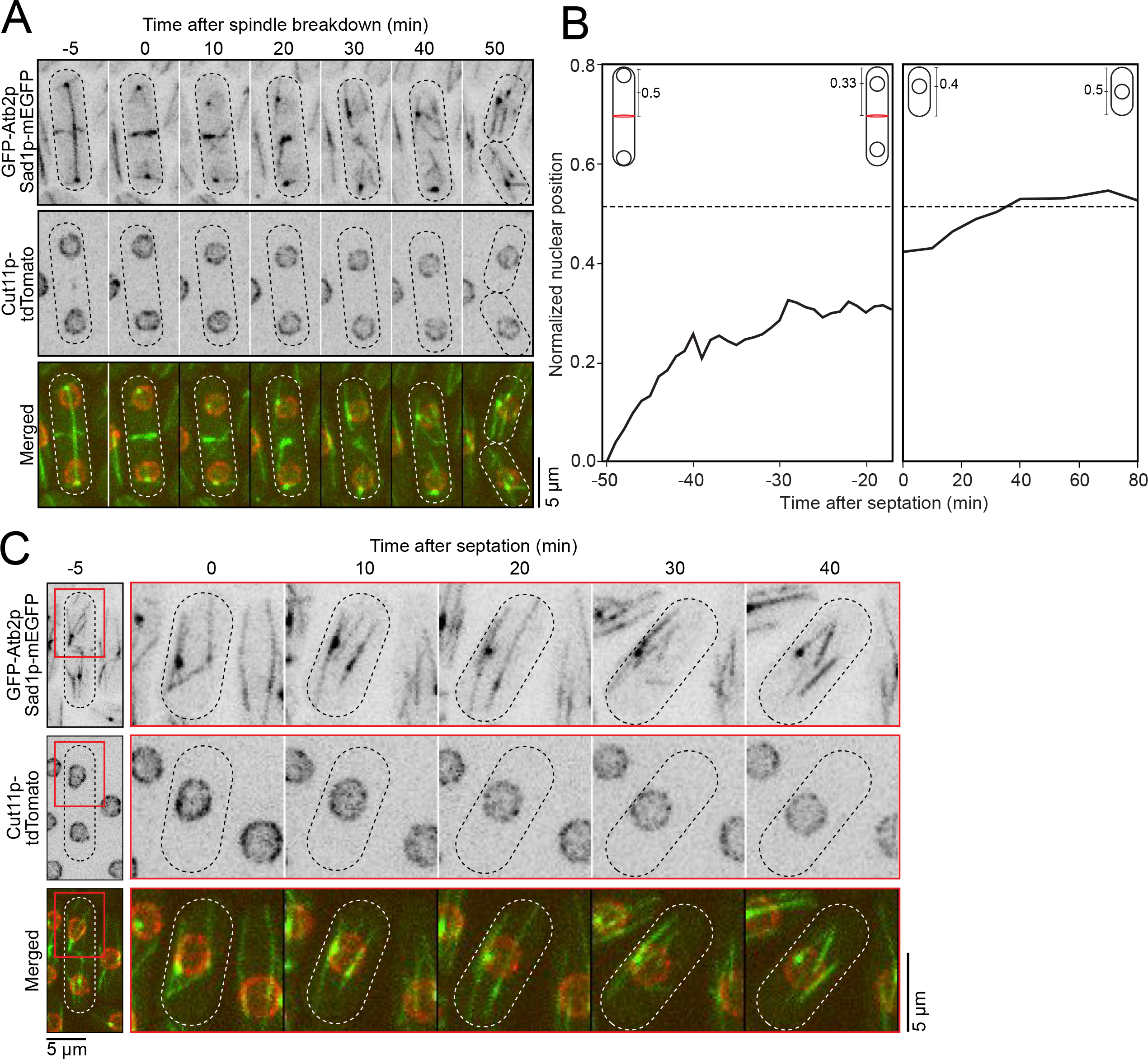
Nucleus is off center from spindle disassembly until ~40 minutes after septation. **A**. Time series of micrographs in a wild-type cell showing nuclear position from spindle disassembly until septation. **B**. Plot of the nuclear position of wild-type cells normalized for the cell length. 0, closest distance to old cell end. 1, closest distance to new cell end. Solid line, average. Dashed horizontal line, normalized position 0.5. n=12-22 cells. **C**. Time series of micrographs of a wild-type cell after septation. Red box, selected region of interest. All micrographs, inverted grayscale for single color, merged for two colors and dotted lines are cell outlines.

These two phases suggested that distinct mechanisms reposition the nucleus after spindle disassembly. We investigated whether microtubules repositioned the nucleus after spindle disassembly by treating cells with 50 μg/mL methyl benzimidazole carbamate (MBC) to depolymerize the microtubule network and then rapidly processed the cell samples for timelapse imaging (Tran *et al*., 2000; Tran *et al*., 2001; Sawin and Snaith, 2004; Castagnetti *et al*., 2010). At the start of imaging, <7 min after MBC addition, interphase microtubules were already depolymerized. Spindles and the PAA microtubules depolymerized more slowly with some PAA microtubules visible at the contractile ring throughout the acquisition (Figure 2A). All nuclear motions, both directed and oscillatory, in cells at any stage of the cell cycle ended upon microtubule depolymerization supporting that microtubules reposition the nucleus after spindle disassembly (Figure 2A and B).

**Figure 2.**
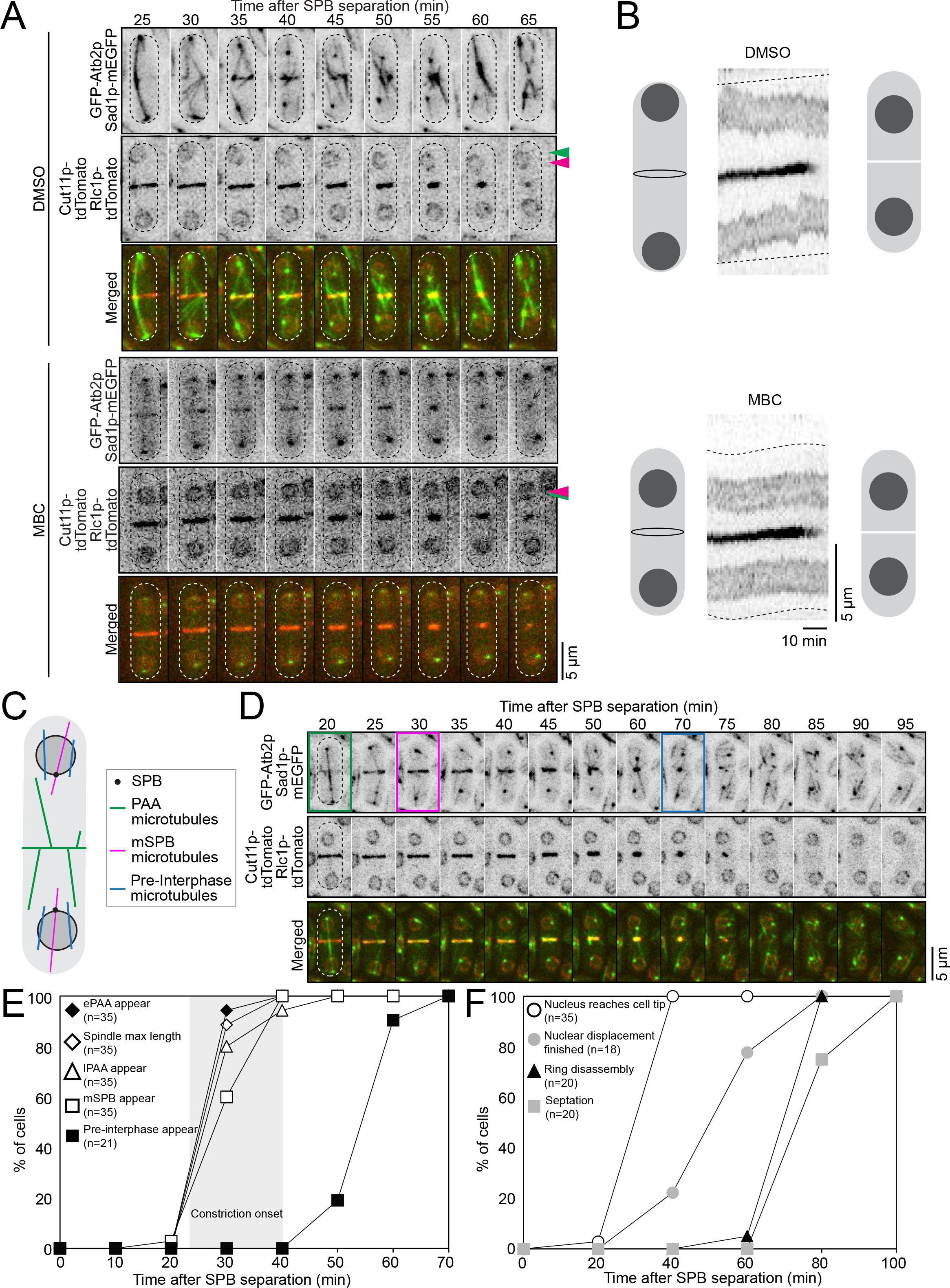
Microtubules position the nucleus after anaphase. **A**. Time series of micrographs of wild-type cell treated with either DMSO or MBC. Green arrowhead, nuclear position at the time of spindle disassembly. Magenta arrowhead, stable nuclear position after displacement. Some ePAA microtubule signal is detectable during the entire acquisition (black arrow). Time 0 is SPB separation for DMSO-treated cells, and MBC-treated cells were aligned to DMSO-treated cells based on the size of their contractile ring. **B**. Kymographs (center) of maximum projections of optical sections of a cell taken at 1-min intervals and diagrams (left, right) for orientation. Dotted lines track the cell ends over time. **C**. Diagram showing the three microtubule networks that polymerize from spindle disassembly until septation. **D**. Time series of micrographs of a wild-type cell. Colored boxes highlight frame with appearance of each of the three networks of microtubules in this example. Color code same as in C. **E, F**. Outcomes plot showing the timing of microtubule network events (E) or nuclear positioning events (F). Number of cells analyzed in parentheses. All micrographs, inverted grayscale for single color, merged for two colors and dotted lines are cell outlines.

Three distinct networks of microtubules polymerize after spindle disassembly: the PAA, the mSPB, and interphase-like microtubules (Figure 2C) (Hagan and Hyams, 1988; Hagan and Yanagida, 1997; Samejima *et al*., 2010). To determine which of these networks contributed to nuclear repositioning, we measured their timing of appearance using timelapse micrographs (Figure 2D and E). The PAA microtubules form a complex network with the equatorial PAA (ePAA) microtubules organized in the plane of the contractile ring and the longitudinal PAA (lPAA) microtubules extending from the contractile ring toward both ends of the cell (Figure 2C and D). The ePAA microtubules appeared at the contractile ring shortly before the spindle disassembled at 25 ± 3 min (mean ± standard deviation, SPB separation = 0 min.) (Figure 2E). The lPAA microtubules were first detected at 29 ± 5 min. The PAA microtubule network remained until the contractile ring disassembled, ~65 min after SPB separation. The mSPB microtubules (Samejima *et al*., 2010) appeared at 30 ± 4 min. This bundle of microtubules appeared to persist beyond septation into the newborn cell and presumably became the interphase microtubule bundle connected to the SPB. Finally, interphase-like microtubules that are not associated with the SPBs appeared at 55 ± 6 min, after the nucleus reached its destination. These microtubules also appeared to persist beyond cytokinesis and possibly became interphase microtubules.

We measured the timing of the nuclear movements from spindle disassembly until septation to correlate the appearance of these microtubules with nuclear repositioning (Figure 2F). In wild-type cells, the nucleus reached the cell ends at 27 ± 4 min, roughly coinciding with the onset of constriction of the contractile ring and the moment when the ends of the elongating spindle reached the cell ends at 25 ± 3 min. The spindle then bowed and disassembled at 29 ± 4 min. Over the following 20 ± 11 min, the nucleus moved back toward the contractile ring reaching its destination at 50 ± 12 min where it oscillated slightly with no further gain in displacement toward the contractile ring until septation. The contractile ring disassembled ~15 min later at 66 ± 4 min and septation occurred at 77 ± 8 min. Both the mSPB and PAA microtubules are present from ~29 min to ~50 min after SPB separation during nuclear repositioning suggesting that they may cooperate to reposition the nucleus from the end of anaphase until septation. Based on our timing measurements, the interphase-like microtubules were not present during the repositioning of the nucleus and were not investigated further.

After septation, bundles of interphase microtubules are present in the cell suggesting that they may be poised to promptly center the nucleus in the newborn cells (Figure 2D). At that moment, the nucleus is only ~1 μm away from the cell center yet this final centering occurs very slowly. The slow centering and the displacement profile during this final phase may indicate that additional factors impact the microtubule-based centering mechanism.

### The mSPB and PAA microtubules compete to position the nucleus

The gamma tubulin complex linker protein Mto1p, a component of MTOCs, is involved in the polymerization of different microtubules throughout the cell cycle (Sawin et al., 2004; Venkatram et al., 2004; Zimmerman and Chang, 2005; Samejima et al., 2008). Deletion of the *mto1* gene (*Δmto1*) causes multiple microtubule defects including the loss mSPB microtubules and PAA microtubules (Venkatram *et al*., 2004; Zimmerman and Chang, 2005; Samejima *et al*., 2010). We leveraged specific mutations in *mto1* that disrupt either the PAA or mSPB microtubules to determine their distinct role in repositioning the nucleus. The *mto1-427* allele disrupts PAA microtubules whereas a 30 amino acid deletion at the C-terminus of Mto1p, the *mto1(1-1085)* allele, disrupts astral and mSPB microtubules specifically (Samejima *et al*., 2010). In addition, cells lacking the myosin-II Myp2p (Δ*myp2* cells) also lack PAA microtubules as Myp2p is required for the recruitment of Mto1p to the contractile ring (Samejima *et al*., 2010).

We acquired timelapse micrographs of *Δmto1*, *mto1-427*, *Δmyp2* or *mto1(1-1085)* mutant cells and measured the destination of the nucleus as the position of the center of the nucleus when it reached its maximal distance from the cell end before the end of cytokinesis (Figure 3A). We normalized the values according to cell length to account for mild variations in cell length across the different genotypes. A destination of 0 represents the position of a nucleus abutting the end of the cell and 1, abutting the contractile ring (Figure 3B). In wild-type cells, the nucleus traveled until it reached a destination of 0.33 ± 0.10. In *mto1-427* (and *Δmyp2*) cells that lack PAA microtubules, the nucleus traveled closer to the contractile ring than in wild-type cells, reaching a destination of 0.42 ± 0.13 (0.44 ± 0.08 for *Δmyp2*). When ring constriction is delayed (*Δmyp2 myo2-E1* cells) in cells that lacked PAA microtubules, the nucleus just crossed the center and reached 0.53 ± 0.14 (Mana-Capelli *et al*., 2012; Laplante *et al*., 2015). In contrast, cells that lack mSPB microtubules, *mto1(1-1085)* cells, the nucleus remained close to the end of the cell with a destination of 0.13 ± 0.09. In those cells, a minimal drift of the nucleus away from the cell end occurred upon spindle disassembly and may result from the relief of the pushing forces exerted by the spindle. Interphase-like microtubules were present in *mto1(1-1085)* cells, supporting that these microtubules are insufficient for repositioning the nucleus after anaphase, consistent with their timing of appearance (Figure 2E). In *Δmto1* cells, cells that lack both PAA and mSPB microtubules, the nucleus remained close to the cell end with a destination of 0.15 ± 0.09, comparable to the destination measured in *mto1(1-1085)* cells. Our observations suggest that the mSPB microtubules move the nucleus away from the cell end while the PAA microtubules limit that movement.

**Figure 3.**
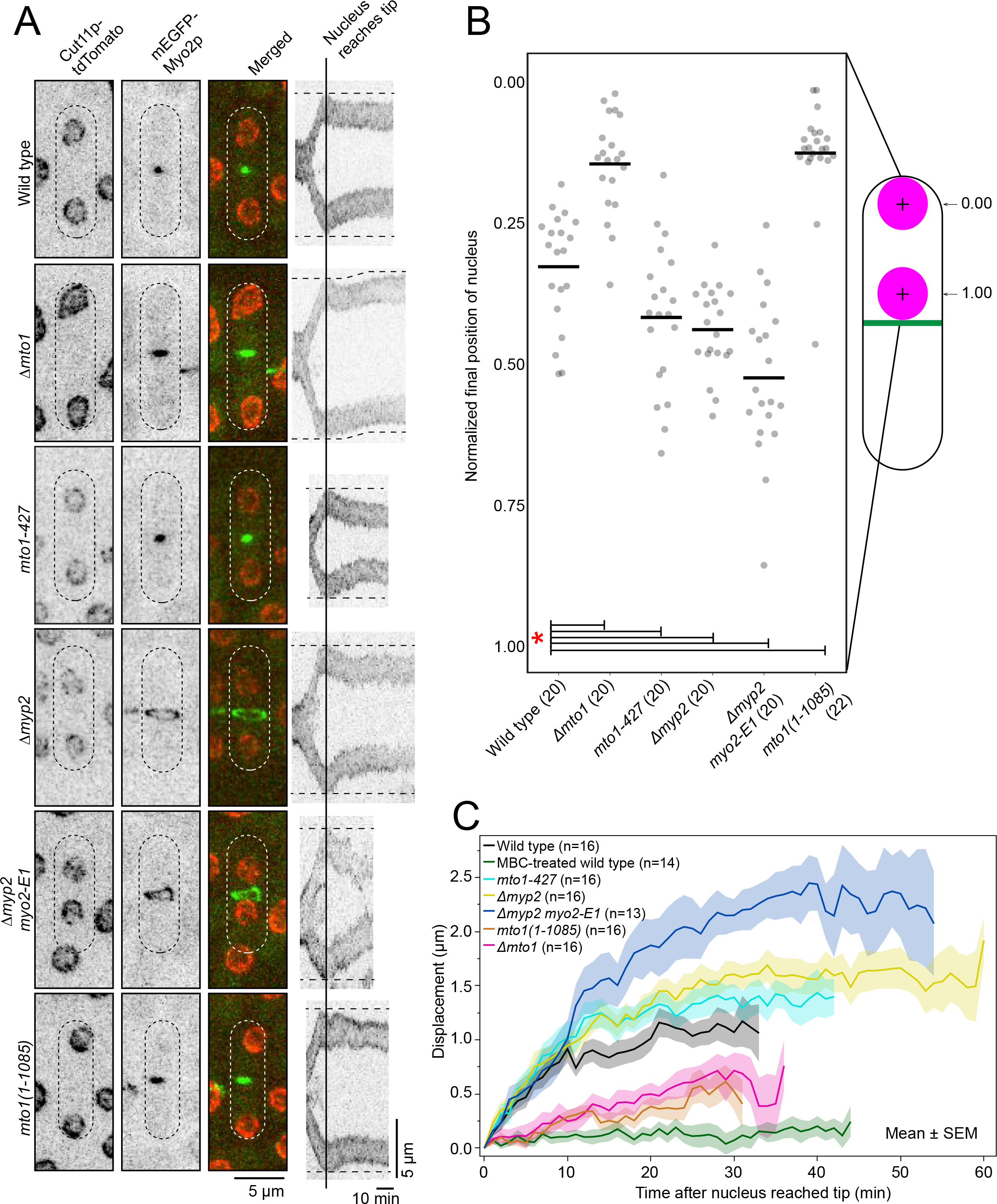
PAA and mSPB microtubules position the nucleus after anaphase until septation. **A**. Confocal micrographs of the final position of the nucleus before ring disassembly (left). Kymographs of maximum projections of optical sections of cells containing Cut11p-tdTomato taken at 1-min intervals aligned at the time when the nucleus reached the cell end (right; dotted horizontal lines, cell ends). **B**. Swarm plot of the nuclear position normalized for cell length. 0.00, closest distance to old cell end. 1.00, closest distance to new cell end. Asterisk marks a significant difference between the indicated comparisons at p < 0.05, Student’s t-test. **C**. Graph of the displacement of the nucleus. Time 0, time when nucleus reaches the ends of the cell by the action of the spindle. Tracking of the nucleus ended when the contractile ring was fully disassembled. Number of cells analyzed in parentheses. All micrographs, inverted grayscale for single color, merged for two colors and dotted lines are cell outlines.

To determine how the two networks of microtubules contribute to the motion of the nucleus, we tracked the displacement of the nucleus every minute from the time it reached the end of the cell until the contractile ring disassembled (Figure 3C and Supplemental Figure 2). In wild-type cells, the mean displacement of the nucleus showed an initial rapid phase during the first ~10 minutes followed by a slower more variable phase. During the rapid phase, the nucleus moved at a mean velocity of 0.09 ± 0.05 μm/min and then dropped to 0.005 ± 0.02 μm/min during the slower phase of displacement. The profile of displacement of the nucleus in cells that lacked mSPB microtubules (*mto1(1-1085), Δmto1* and cells treated with MBC) showed a single slow phase of displacement. In those cells, the nucleus displaced slowly from spindle breakdown until the disassembly of the contractile ring (0.05 ± 0.02 μm/min for *mto1(1-1085)*, 0.06 ± 0.03 μm/min for *Δmto1* and 0.04 ± 0.02 μm/min for MBC treated cells). These results suggest that the initial rapid phase of displacement observed in wild-type cells depends on the actions of the mSPB microtubules (Figure 3C and Supplemental Figure 2). The nuclear displacement traces for cells that lack PAA microtubules (*mto1-427*, *Δmyp2* and *Δmyp2 myo2-E1*) followed a similar profile as that of wild-type cells with an initial rapid phase followed by a slow phase (Figure 3C). The velocity of the nuclear movement during the initial phase of displacement in *mto1-427*, *Δmyp2* and *Δmyp2 myo2-E1* was comparable to that of wild-type (0.10 ± 0.06 μm/min for *mto1-427*, 0.09 ± 0.05 μm/min for *Δmyp2* and 0.11 ± 0.05 μm/min for *Δmyp2 myo2-E1*) (Supplemental Figure 2). The nuclear velocity of the second phase of displacement was similar to wild type in *mto1-427* (0.01 ± 0.02 μm/min, p = 0.23), and slightly faster in both Δ*myp2* (0.02 ± 0.03 μm/min, p < 0.05) and in *Δmyp2 myo2-E1* cells (0.03 ± 0.02 μm/min, p < 0.001) (Supplemental Figure 2). The transition between the initial rapid phase and slow phase was delayed by ~5-10 min in cells that lacked PAA microtubules. This delayed transition suggests that the PAA microtubules intercept the nucleus ~10 min after the onset of nuclear motion thus ending the initial fast phase of motion.

### mSPB microtubules push the nucleus away from the cell ends

To understand how mSPB microtubules displace the nucleus toward the division plane, we characterized their organization and dynamics. The entire mSBP microtubule is composed of a proximal and distal bundle (Figure 4A). We measured the fluorescence intensity of the mSPB bundles labeled with GFP-Atb2p, compared it to that of the interphase microtubule bundles, expected to be composed of 4 microtubules, and found that mSPB microtubules are bundles of 3-4 microtubules near the SPB tapering down to 1-2 microtubules near their +tips (Tran *et al*., 2001; Loiodice *et al*., 2005; Daga *et al*., 2006). The average length of the proximal bundle gradually increased during nuclear displacement until it reached a maximal length of 2.9 ± 0.7 μm (Figure 4B). The distal bundle was consistently shorter than the proximal with a maximal length of 1.6 ± 0.7 μm. Both proximal and distal bundles showed dynamic instability characterized by periods of growth followed by rapid depolymerization typical of microtubules. However, the proximal bundle appeared more stable and contacted the ends of the cell for 2.6 ± 1.3 min (n = 20) before shrinking. In contrast, the distal bundle shrunk more frequently and did not contact any obvious structure. We concluded from these results that the mSPB microtubules may push rather than pull the nucleus toward the division plane.

**Figure 4.**
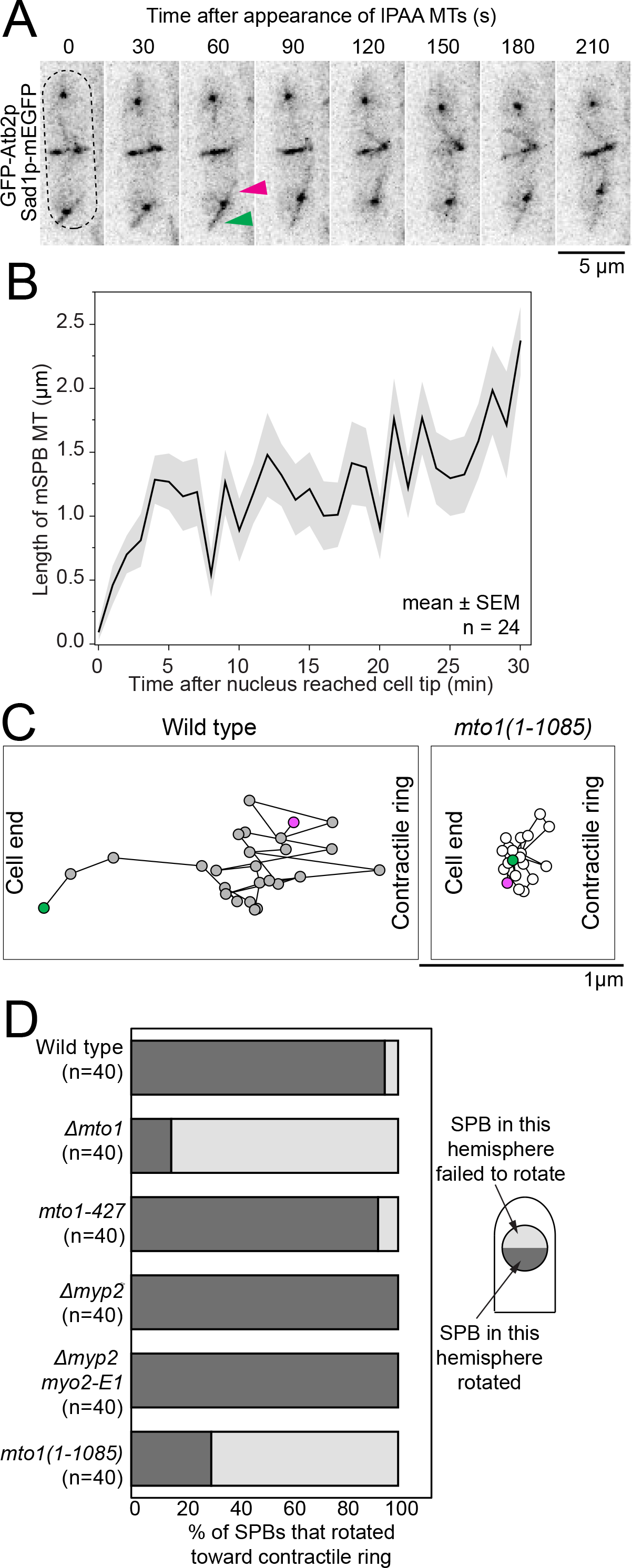
Elongation of mSPB microtubules against the cell end rotates and pushes the nucleus. **A**. Time series of micrographs of a wild-type cell showing mSPB microtubule dynamics. Green arrowhead, proximal mSPB bundle. Magenta arrow, distal mSPB bundle. **B**. Plot of the length of the proximal mSPB bundle over time in wild-type cells. **C**. Representative traces of SPB position taken every 1 min in wild-type and *mto1(1-1085)* cells from the time the nucleus reached the end of the cell until ePAA disappearance. Green circle, starting position. Magenta circle, ending position. **D**. Stacked bar chart of the percentage of SPBs that rotated to face the contractile ring before septation. Diagram of scoring of SPB rotation (right). Micrographs, inverted grayscale for single color and dotted line is cell outline.

We expected such pushing motion of the mSPB microtubules at the SPB to rotate the nucleus and the rotation to be detectable in the position of the SPBs embedded in the nuclear envelope. At the start of nuclear displacement shortly after the spindle disassembled, the SPB faced the end of the cell likely due to the pushing action of the spindle on the SPBs (Figure 4C and, D and Supplemental Figure 3) (McCully and Robinow, 1971; Tanaka and Kanbe, 1986; Ding *et al*., 1993). Concurrent with the appearance and elongation of the mSPB microtubules, the SPBs rotated toward the division plane in wild-type cells. In the absence of mSPB microtubules, in *mto1(1-1085)* and *Δmto1* cells, the SPBs did not rotate and remained facing the cell end (Figure 4C, D and Supplemental Figure 3). Our observations suggest that mSPB microtubules pushed against the cell end and rotated the nucleus causing the reorientation of the SPBs. Consistent with this conclusion, we often observed the deformation of the nuclear envelope in response to the mSPB microtubules pushing against the SPB. This effect was also observed when interphase microtubules contact and push against end of the cell transferring the pushing force to the SPB thus deforming the nuclear envelope as the nucleus moves away from the cell end (Tran *et al*., 2001; Daga and Chang, 2005).

### The PAA microtubules basket the nucleus and limit its movement toward the contractile ring

The basket shape of the PAA microtubule network suggests that they may physically intercept the incoming nucleus (Figure 5A and B). The lPAA microtubules are very dynamic with constant elongation and shortening back to the ePAA microtubules that rearrange the entire network on a timescale of minutes (Figure 5C). We quantified the contacts between the lPAA microtubules and the nucleus in cells expressing GFP-Atb2p or the EB1 homolog Mal3p-mEGFP and Cut11p-tdTomato (Figure 5D and E). We scored the colocalization of the two markers as contacts between the microtubule and the nucleus. The lPAA microtubules made frequent contacts with the nucleus either with their +tip or along their side (Figure 5D). The number of contacts between the lPAA microtubules and the nuclear envelope ranged from 1 to 3 per camera frame and each contact lasted from ~20 to 50 s (Figure 5F and G). Therefore, the PAA microtubules likely make physically contacts with the nucleus possibly causing the previously measured transition between the fast and slow nuclear displacement phases (Figure 3C). Consistent with this interpretation, the nucleus abruptly moved closer to the division plane when the contractile ring and PAA microtubule network disassembled (Figure 1B).

**Figure 5.**
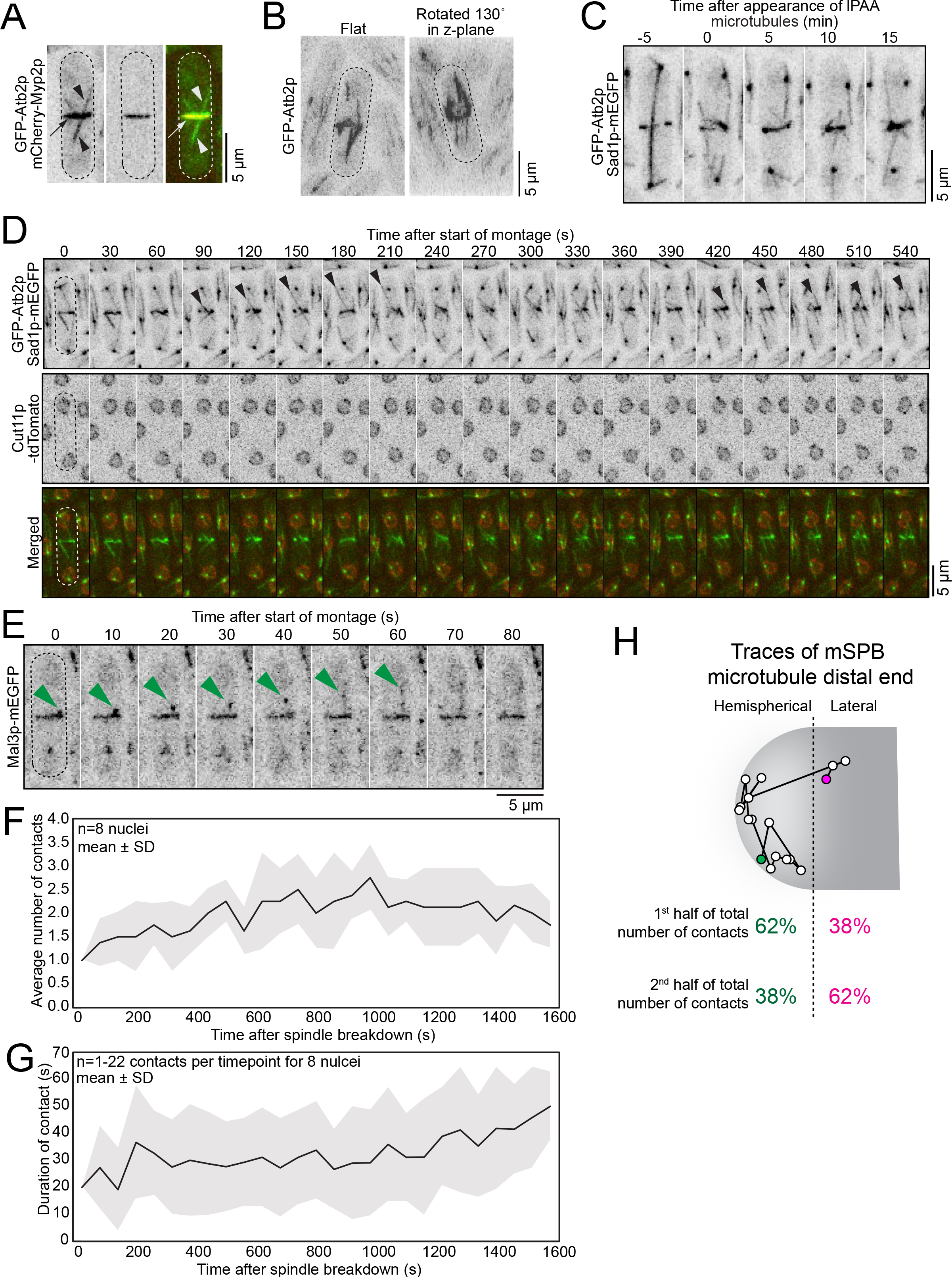
PAA microtubules position the nucleus through direct, dynamic contact. **A**. Confocal micrographs showing ePAA (arrow) and lPAA microtubules (arrowheads) in a wild-type cell. **B**. Flat (left) and rotated 130° (right) micrographs of the PAA microtubules. **C**. Time series of micrographs showing the dyamics of PAA microtubules in a wild-type cell. **D**. Time series of micrographs of strains showing PAA microtubules and nuclei in a wild-type cell. Arrowheads, contacts between lPAA and nucleus. **E**. Time series of micrographs showing +tips of lPAA microtubule (green arrowheads) extending toward nucleus. **F, G** Plots of the number of contacts by the lPAA microtubules with the nuclear membrane (F) and the duration of these contacts (G) over time. **H**. Representative trace of mSPB microtubule position taken every 1-min in a wild-type cell from the time of mSPB microtubule appearance to ring disassembly. Green circle, start position. Magenta circle, end position (Top). Percentage of location of contacts of the proximal end of the mSPB microtubule during the first and second half of the total number of contacts (Bottom). n=20 mSPB microtubules. All micrographs, inverted grayscale for single color, merged for two colors and dotted lines are cell outlines.

In the absence of PAA microtubules, the nucleus moved closer to the contractile ring suggesting that the proximal mSPB bundles may grow longer in cells lacking PAA microtubules (Figure 3B and C). However, in *mto1-427* cells, the proximal mSPB bundles reached a maximum length of 2.9 ± 0.8 μm (n = 20) while the distal mSPB bundle reached a length of 1.4 ± 0.4 μm, comparable to the lengths measured in wild-type cells. We mapped the contacts points of the +tip of the proximal mSPB microtubules along the inner cell surface from the time of mSPB appearance until ring disassembly. The proximal mSPB bundle contacted the end of the cell transiently in a touch-and-retract fashion. For the first half of the total number of measured contacts (~the first 15 minutes of repositioning), 62% of contacts were made within the hemispherical end of the cell in wild-type cells (n = 20 mSPB microtubules; Figure 5H). During the second half of all contacts (~the last 15 minutes of repositioning), 62% of the contacts were made with the lateral sides of the cell. Therefore, in the absence of PAA microtubules, the proximal mSPB microtubules contact the cell surface progressively closer to the division plane to push the nucleus toward the division plane. Such progressive stepping may explain how short microtubules can push the nucleus to distances that exceed their length. The actions of the PAA microtubules may then restrict the contacts of the mSPB microtubule to the cell ends and prevent their progressive stepping toward the contractile ring.

### Position of the nucleus within the cell predicts the velocity of nuclear displacement

Interphase microtubules can reposition an artificially displaced nucleus with a peak displacement velocity ranging from 0.2-0.9 μm/min (Tran *et al*., 2001; Daga and Chang, 2005; Tolic-Norrelykke *et al*., 2005). We therefore expected the interphase microtubules, present in the newborn cell at the time of septation, to rapidly center the nucleus within 5 minutes after septation. In contrast, the nucleus slowly reached the cell center over the first ~40 min after septation (Figure 1B and C, and Supplemental Figure 4A). The displacement profile of the nucleus in a wild-type cell showed an initial static phase of ~10 min followed by a slow displacement of ~0.01 μm/min (mean velocity) until the nucleus reached a final stable position ~40 min after septation (Figure 1B and Supplemental Figure 4A). The absence of net nuclear movement in the first 10 min after septation suggests that although microtubules are present at that time, their dynamics do not result in any significant nuclear movements.

To understand the molecular mechanisms responsible for this slow displacement we modified a previous model of nuclear displacement in fission yeast cells by Foethke *et al*. (Supplemental Table 2) (Foethke *et al*., 2009). We first simulated the repositioning of an off-center nucleus located at the cell end in a 12.4 μm long cell, the same cell length studied by Daga *et al*. The simulations effectively repositioned the nucleus to the cell center at a peak displacement velocity, the initial velocity of the displacement, of 0.34 μm/min, in agreement with the range measured by Daga *et al*. (Supplemental Video 1) (Daga *et al*., 2006). We measured a mean velocity of the entire displacement of 0.08 μm/min, nearly 10-fold faster than the nuclear displacement velocity we measured in newborn cells.

To understand the factors that influence nuclear displacement velocity, we observed the video output from simulations of nuclear displacement in short and long cells that have a nucleus positioned against the end of the cell. Videos showed dynamic microtubules growing nearly parallel to the long axis of the cell from sites anchored at the nuclear envelope (Supplemental Video 1). Cycles of microtubule polymerization followed by sudden catastrophe captured the dynamic instability of microtubules. Microtubules polymerized until they reached the closest cell end, bowed slightly while exerting pushing forces that moved the nucleus away from that cell end, and rapidly underwent catastrophe. We refer to the distance between the nucleus and the proximal end of the cell as the proximal (P) distance and the distance between the nucleus and the distal end of the cell as the distal (D) distance. In a newborn wild-type cell, the proximal end is the old cell end, and the distal end refers to the new cell end created by septation. In videos of long cells (12.4 μm) with a nucleus located at the cell end, the proximal microtubules visibly reached the proximal cell end before distal microtubules could reach the distal end of the cell. Therefore, the proximal microtubules had a pushing advantage over the distal microtubules at every round of polymerization and catastrophe. From these observations, we hypothesized that the P/D ratio influenced the velocity of the nuclear displacement. We ran simulations with nuclei positioned across a range of physiologically relevant P/D ratios for wild-type cells of different lengths. Nuclear velocity decreased with increasing P/D ratio supporting that the greater the difference between the P and the D distances the faster the nucleus repositioned to the cell center (Figure 6B).

**Figure 6.**
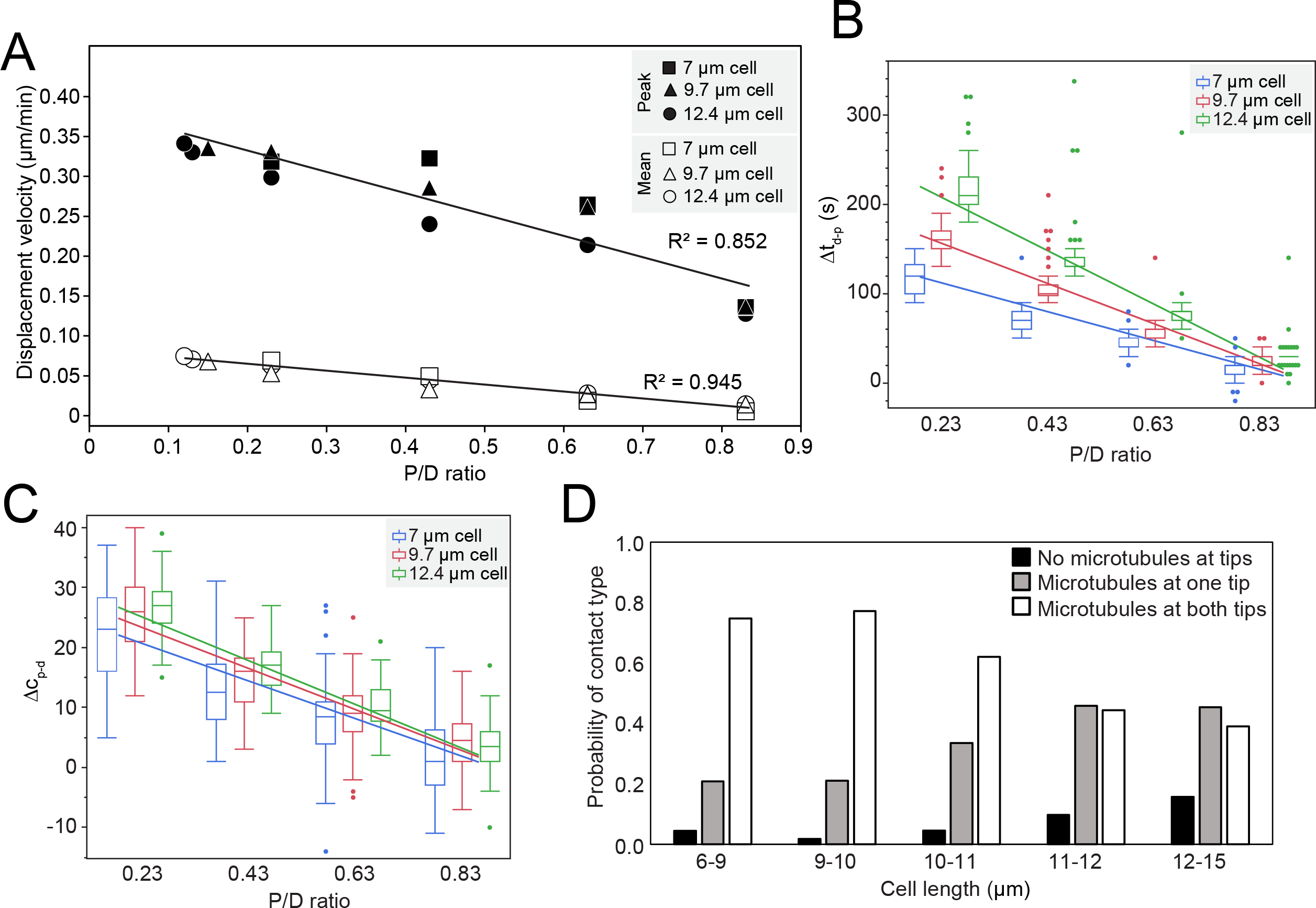
P/D ratio determines nuclear displacement velocity. **A**. Correlation plot between the nuclear displacement velocity and the P/D ratios simulated for cells of different lengths. Line, linear fit. n=50 simulations per cell length and P/D ratio combination. **B, C**. Correlation plots between the matched P/D ratios of cells of different lengths and either Δt_d-p_ (C) or Δc_p-d_ (D). Line, linear fit. n=50 simulations per cell length and P/D ratio combination. **D**. Histogram of the probability of microtubules simultaneously touching at both cell ends, touching one cell end or not touching either cell end in cells of different lengths.

This relationship predicts that the proximal microtubules have a pushing advantage over the distal microtubules. We extracted information about the dynamics of microtubules +tips from simulations and measured the additional time required for the distal microtubules to reach the distal cell end over the time required for proximal microtubules to reach the proximal cell end (Δt_d-p_) at the start of the simulations. We also measured the additional number of microtubule contacts at the proximal cell end over those at the distal cell end (Δc_p-d_) for the first 20 min of the simulation. We used Δt_d-p_ and Δc_p-d_ as quantitative values for measuring asymmetric pushing advantage. We found that both Δt_d-p_ and Δc_p-d_ increased with decreasing P/D ratios (Figure 6B and C). In addition, both Δt_d-p_ and Δc_p-d_ increased with increasing cell length. Our simulations support that proximal microtubules have a greater pushing advantage when the nucleus is positioned farther off center *i.e*. at a smaller P/D ratio. Finally, we determined whether the P/D ratio could recapitulate the measured nuclear displacement velocity measured in newborn cells. Simulations of a 7-μm cell with a nucleus located at a relative nuclear position of 0.4 (P/D ratio of 0.83), comparable to a wild-type newborn cell, resulted in a mean nuclear displacement velocity of 0.01 μm/min, the same velocity measured experimentally. Similarly, a P/D ratio of 0.44 in a 7-μm cell, similar to *mto1-1085* cells predicted a mean displacement velocity of 0.04 μm/min, comparable to the experimental value of 0.05 μm/min. Our simulations support that nuclear displacement velocity increases when the proximal microtubules exert pushing forces that are unopposed by the actions of the distal microtubules while nuclear velocity decreases with increasing simultaneous pushing against both cell ends.

Our simulations predicted a high frequency of simultaneous microtubule contacts at both cell ends in newborn cells. We experimentally measured microtubule contacts with the ends of the cell over time from cell birth (~7-μm cell) until the disassembly of the interphase microtubules ahead of spindle assembly (~14-μm cell) while keeping track of cell size (Figure 6D). From those measurements, we calculated the probability of microtubule contacts at both cell ends simultaneously, at one cell end only and no contact at either end. In cells ranging from 6 to 10 μm in length, the probability of simultaneous contacts was 0.76 while the probability of single contact was 0.2 and the probability of no contact was <0.1. In longer cells ranging from 11 to 15 μm, the probability of simultaneous and single contacts was similar at ~0.4 each. Interestingly, the probability of simultaneous contacts drops when cells reach ~10 μm in length. Therefore, as predicted from our simulations, we found that the probability of microtubules to contact both ends of the cell simultaneously is highest in short cells and lowest in longer cells.

Our results suggest that the difference between the proximal and the distal distances influences nuclear velocity. Smaller P/D ratios result in faster nuclear movements presumably due to net asymmetric pushing against the proximal cell end, resulting in a greater velocity. In contrast, larger P/D ratios result in frequent simultaneous pushing against both cell ends. These inefficient pushing actions result in slow nuclear displacement velocities. Our experimental data suggest that in short cells, proximal and distal microtubules simultaneously contact their respective cell end with high frequency likely resulting in a “tie” in the pushing forces that slows down nuclear displacement.

### Cell growth is required to recapitulate the nuclear displacement profile in the newborn cell

Simulations based on microtubule dynamics only failed to reproduce the nuclear displacement profile measured in live cells (Figure 7A). The profile of nuclear displacement in newborn cells showed an initial 10 min static phase immediately after septation followed by a slow displacement during which the nucleus moves to its final stable destination located just beyond the cell center (normalized position ~0.54) ~40 min after septation. In contrast, nuclear displacement profile from simulations showed no initial plateau and a premature stabilization of the nucleus at the cell center (normalized position ~0.50) ~20 min after septation. These results suggested that additional factors besides the previously described interphase microtubule dynamics influence nuclear displacement in live cells.

**Figure 7.**
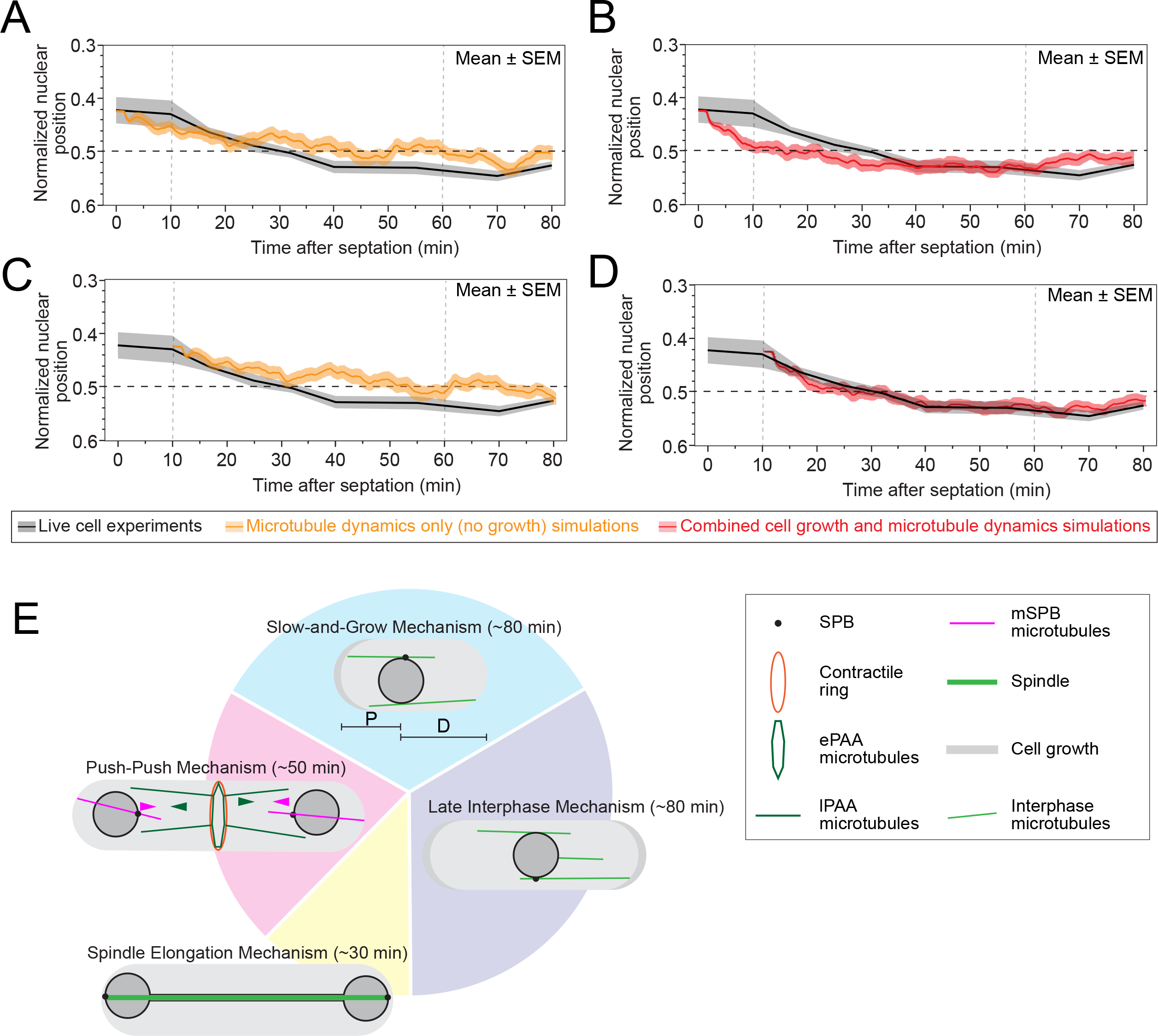
Cell growth cooperates with interphase microtubules dynamics to center the nucleus after septation. **A, B**. Plots of normalized nuclear position in 7-μm cells with a nucleus positioned at 0.83 P/D ratio in simulations with microtubule dynamics only (A, orange) and simulations with cell growth and microtubule dynamics combined (B, red). Mean of 50 simulations for each A and B. Experimentally measured normalized nuclear position in wild-type cell (same as Supplemental Figure 4A). **C, D**. Same plots as in A and B aligned at the start of nuclear displacement in the experimental data 10 minutes after septation. Horizontal black dashed line, center of cell. Vertical grey dashed lines, time of OETO and NETO respectively. **E**. Model of nuclear positioning mechanisms and their expected duration across the cell cycle. In newborn cells, a slow-and-grow mechanism slowly centers the nucleus. Once cells reach 9.5-10 μm in length, the late interphase nuclear positioning mechanism maintain/correct the center position of the nucleus until anaphase. During anaphase, the spindle pushes the two daughter nuclei to opposite ends of the cells. From spindle disassembly until septation, a push-push mechanism repositions the nucleus from the cell end until a third of the distance from the cell end. Time in parentheses are approximate durations for each mechanism based on a 4-hour cell cycle.

Although cell growth occurs at a timescale negligible compared to rapid microtubule dynamics, nuclear displacement in short cells with a nucleus positioned at a high P/D ratio occurs on a timescale similar to cell growth. We thus considered that cell growth may contribute to the nuclear displacement profile in newborn cells. We measured the profile of cell growth in the first 80 min after septation. Mitchison and Nurse detected an early delay in the growth of the old end (Old End TakeOff, OETO) of some cells after septation (Mitchison and Nurse, 1985). We measured the profile of cell growth by tracking the appearance of For3-3xGFP, a component of the polarisome, or actin patches, marked with GFP-CHD, in newborn wild-type cells (Feierbach *et al*., 2004; Martin and Chang, 2005; Martin *et al*., 2005). We first detected For3-3xGFP at the old cell end 12 ± 5 min after septation and at the new end 57 ± 7 min (New End TakeOff, NETO) after septation (Supplemental Figure 4B and C). We measured the growth rate of a newborn cell from 10 to 40 min after septation to be 0.024 μm/min, similar to the previously measured growth rate (Mitchison and Nurse, 1985). Using the measured asymmetric growth profile and rate, we built a kinematic model with no microtubule dynamics and a static nucleus inside an asymmetrically growing cell (Supplemental Figure 4D). The kinematic or growth-only model reproduced the overall shape of the experimental profile of nuclear displacement including the initial plateau and the final stable position of the nucleus beyond the cell center, suggesting that cell growth may be a key factor in the nuclear displacement profile.

The kinematic/growth-only model assumes an immobile nucleus inside a growing cell with no microtubule dynamics, a situation that cannot be experimentally reproduced as depolymerization of microtubules to immobilize the nucleus also impacts cell polarity and cell growth (Verde *et al*., 1995; Browning *et al*., 2000; Brunner and Nurse, 2000; Snaith and Sawin, 2003; Feierbach *et al*., 2004; Snaith *et al*., 2005). To test the impact of cell growth on nuclear displacement, we modified the simulation platform that only considered microtubule dynamics to allow the cell to grow according to our measured growth profile thus creating a growth and microtubule dynamics simulation platform. We simulated nuclear displacement in a 7-μm wild-type cell growing asymmetrically from OETO 10 min after septation and then symmetrically starting at NETO 60 min after septation. Growth rate at each cell end was set to the measured rate of 0.024 μm/min. The nucleus was initially positioned at a P/D ratio of 0.83 while all other parameters and variables remained unchanged from our previous simulations (Supplemental Video 2). Nuclear displacement from the growth and microtubule dynamics simulations improved the overall fit with the experimental displacement profile and recapitulated the destination of the nucleus to a normalized position of ~0.54 (Figure 7B). Adding cell growth did not change the mean nuclear displacement velocity of 0.01 μm/min obtained in our previous simulations without cell growth. Although the fit improved, the simulations with growth and microtubule dynamics still did not capture the initial 10 min plateau in nuclear position (p < 0.001, Tukey’s HSD). The molecular reasons for the initial 10 min plateau remain unknown but our results suggest that microtubules may be incompetent to move the nucleus or that the nucleus may not be mobile during that time.

As the simulations reproduced the profile of the active displacement of the nucleus and recapitulated the nuclear displacement velocity measured in live cells, we focused on the section of the profile starting 10 min after septation. We thus aligned the start of our simulations to the 10 min mark of the experimentally measured nuclear displacement in live cells. This realignment resulted in a better agreement between the simulated data with growth and the live cell displacement profile. The simulations with microtubule dynamics only but with no cell growth still showed poor agreement with the experimentally measured nuclear displacement profile (Figure 7C and D). These results further support that our simulation framework based on cell growth and microtubule dynamics can explain nuclear displacement after the initial 10 min plateau. These simulations showed strong agreement with both the final stable position at 0.54 and the time at which the nucleus reached that position. Together our simulations suggest that in short newborn cells with a nucleus positioned at a high P/D ratio, nuclear displacement slows down or stalls. At such a timescale, cell growth becomes relevant in the overall nuclear displacement.

## DISCUSSION

We explored the mechanism that recenters the nucleus after anaphase. Like many other cell types, fission yeast cells use microtubules to reposition their nucleus. Simulations of a nuclear positioning mechanism based on microtubule dynamics only reproduced oscillatory motions and repositioning of the nucleus in late interphase cells (Tran *et al*., 2001; Daga *et al*., 2006; Foethke *et al*., 2009). This mechanism relies on the specific dynamics and organization of microtubules in mid- to late-interphase cells. The intrinsic properties of microtubules that cause their dynamic instability result in temporary alternating pushing forces from microtubules polymerization against the cell ends. Additional factors not included in the previously described mechanism based on late interphase microtubule dynamics were required to explain how the nucleus recentered after anaphase since the general organization of the microtubule network after anaphase is different from the organization of interphase microtubules. We found that cell growth and microtubule competition conjointly influence nuclear positioning from anaphase until ~40 min after septation covering a large portion of the total duration of a fission yeast cell cycle (Figure 7E).

Our measurements suggest that competition between the mSPB and PAA microtubules drives nuclear displacement in a push-push mechanism from the end of anaphase until septation (Figure 7E). Immediately after spindle disassembly, the mSPB microtubules polymerize until the proximal bundle of mSPB microtubules contact the proximal end of the cell where it exerts pushing forces. The force generated by the mSPB microtubules are transferred to the nucleus to rotate and push the nucleus toward the contractile ring. The PAA microtubule network then intercepts the moving nucleus and stops its progression at a third of the distance from the cell end to the contractile ring.

The basket-like structure of the PAA microtubule network and its attachment to the contractile ring are likely key to provide the counterforce necessary to push against the incoming nucleus. The nature of the connections between the eMTOCs and the contractile ring is not fully understood but relies on the direct or indirect interactions between Mto1p and the myosin II Myp2p (Samejima *et al*., 2010). To better understand the PAA microtubules, we will need to uncover their molecular organization and the nature of their attachment to the bundle of actin filaments of the contractile ring.

It is tempting to posit that the function of the PAA microtubules is to block the nucleus from entering the contractile ring and prevent fatal DNA damage. Although we cannot rule out that the lack of PAA microtubules could result in a damaged nucleus/DNA under untested conditions, the lack of PAA microtubules did not result in nuclei being cut by the contractile ring in our experiments. In the absence of the PAA microtubules, the nucleus never reached the contractile ring before septation and rarely crossed the center of the half-cell compartment before septation. If the DNA is not in danger of being damaged in the absence of PAA microtubules, what is the purpose of such a complex network of microtubules? The PAA microtubules keep the nucleus at third of the distance between the cell end and the contractile ring because that position may be the optimal target position to avoid nuclear-associated material from becoming unevenly distributed after cytokinesis. For example, the type I interphase nodes containing the kinases Cdr1p and Cdr2p, and the Anillin/Mid1p distribute in a broad band around the nucleus before septation (Akamatsu *et al*., 2014). A nucleus positioned closer to the ring may result in the altered distribution of these nodes and the factors responsible for their distribution. As these nodes contain cell cycle regulation factors, their uneven inheritance between daughter cells may be deleterious.

The final position of the nucleus in the absence of PAA microtubule suggests that other factors prevent the nucleus from reaching the contractile ring and getting damaged by its constriction. Even when contractile ring closure was delayed (*Δmyp2 myo2-E1*) giving the nucleus more time to travel toward the ring, the nucleus stopped just beyond the center of the half-cell compartment. Other dense material in the vicinity of the division plane may prevent the nucleus from reaching the contractile ring. The actin network of the contractile ring is a possible candidate that can exert forces against the nucleus and influence its position (Yukawa *et al*., 2021). Alternatively, connections between the nuclear envelope and the plasma membrane, such as the cortical endoplasmic reticulum, may slow down the displacement of the nucleus (Zhang *et al*., 2010). Finally, a local increase in cytoplasmic viscosity in the vicinity of the division plane may impede the motion of the nucleus toward the contractile ring. Any of these putative mechanisms would have the same outcome of impeding the progression of the nucleus toward the contractile ring.

The push-push mechanism leaves the nucleus off center after septation and at that moment the interphase microtubules finalize the centering of the nucleus. Our simulations suggest that nuclear centering is slowed down by microtubule competition. This competition involves microtubules simultaneously pushing against both cell ends resulting in a tie in the pushing forces and an ineffective displacement of the nucleus. Our simulations suggest that a tie in the pushing forces arises when both proximal and distal microtubules are roughly the same length (*i.e*., when the nucleus is positioned at a large P/D ratio). This situation occurs naturally in short newborn cells that have a nucleus slightly off center. Our simulations thus predicted a high probability of microtubules simultaneously contacting both cell ends in newborn cells. As predicted, we experimentally measured a high probability of simultaneous microtubule contact at both cell ends in newborn cells. The probability of simultaneous contacts at both cell ends drops once cells reach a length of 9.5-10 μm. The length of microtubules may be a possible explanation for this transition as short microtubules can sustain a greater amount of stress before buckling than longer microtubules (Mickey and Howard, 1995; Felgner *et al*., 1997). Therefore, a tie in pushing forces in longer cells is less likely as longer microtubules buckle more easily under stress. In cells that are ~10 μm in length, microtubules may reach a length at which the stress exerted by a pushing microtubule exceeds the buckling force of the opposite microtubule causing it to break and catastrophe. Thus, cell growth may be a built-in natural tiebreaker in the pushing forces that occurs when cells that reach ~10 μm in length.

Simulations based on microtubule dynamics alone were insufficient to explain the profile of nuclear displacement measured in live cells. Cell growth occurs at a similar timescales as nuclear positioning in newborn cells and the change in cell size caused by growth can influence nuclear positioning based on the intrinsic properties of microtubules as discussed earlier. Combining cell growth with microtubule dynamics improved the fit between the simulated and experimentally measured profiles of nuclear displacement and recapitulated the position of the nucleus at the end of the displacement. Our work supports a slow-and-grow mechanism of nuclear positioning in newborn cells (Figure 7E). In this model, microtubule competition stalls the displacement of a nucleus positioned at a large P/D ratio. This effect is caused by the frequent simultaneous contacts of microtubules at both ends of the cell resulting in a tie in the pushing forces that reduces the velocity of nuclear displacement. When the nucleus is stalled, cell growth becomes a relevant factor in the profile of nuclear displacement. This slow-and-grow mechanism reproduced the displacement of the nucleus from 10 to 40 min after septation and may be the leading mechanism of nuclear positioning until the cell is ~9.5-10 μm in length. When cells reach that length, ~80 min after septation, nuclear positioning may transition from the slow-and-grow mechanism to the late interphase nuclear positioning mechanism (Figure 7E).

None of the simulations that involved microtubule dynamics replicated the initial 10-minute plateau in the nuclear displacement that we measured in live cells immediately after septation. Instead, simulations showed that microtubule dynamics were sufficient to immediately move the nucleus, even with a high frequency of simultaneous microtubule contacts at both cell ends. These results support the existence of additional factors that prevent nuclear displacement immediately after septation. Microtubule dynamics or the connections between microtubules and the nuclear envelope may be different during those first 10 minutes preventing microtubules to move the nucleus during that period. Therefore, the parameters we used in our simulations, which were measured experimentally in longer interphase cells may need adjustments to recapitulate this nuclear behavior (Foethke *et al*., 2009). Alternatively, the properties of the nucleus or of its cytoplasmic environment may be different during that time period. Those properties may effectively act as a “glue” that prevent microtubule dynamics from centering the nucleus during the first 10 minutes in the newborn cell. An interesting possibility to consider is that the factors that “glue” the nucleus during the first 10 min after septation may be the same ones that prevent the nucleus from getting into the contractile ring before septation. Connections between the nucleus and the plasma membrane or a high local cytoplasmic viscosity may hinder nuclear motions from ~20 min before septation until ~10 min after septation.

Our work suggests that different factors work in tandem with microtubule-based mechanisms to position the nucleus. The intrinsic properties of microtubules can lead to different outcomes in the position of the nucleus depending on the changes in cell size and shape that naturally occur throughout the cell cycle. The mechanical properties of the nucleus can impact its positioning throughout the cell cycle by tuning its response to force producing changes in the cytoskeleton. These factors may be conserved in microtubule-based nuclear positioning mechanisms across different cell types. Given the abundance of knowledge of microtubule dynamics in fission yeast and the ease of genetic manipulations, fission yeast is a prime model organism to uncover unknown factors that govern nuclear displacements. In combination with computational simulations, these powerful toolsets can be utilized to understand the mechanisms of nuclear positioning.

## METHODS

### Strains, Growing Conditions, and Genetic and Cellular Methods

Supplemental Table 1 lists the *S. pombe* strains used in this study. Strains were created using PCR based gene targeting to integrate the constructs into the endogenous locus, except *Pnmt41-GFP-CHD*, which was integrated into the *leu1* locus (Bahler *et al*., 1998). Either *pFA6a-mEGFP-kanMX6* or *PFA6a-tdTomato-kanMX6* were used to create templates to generate C-terminal tagged constructs. The primers had 80 bp of homologous sequence surrounding the integration site (sourced from www.bahlerlab.info/resources/). Except for the *Pnmt81-GFP-atb2* and *Pnmt41-GFP-CHD* constructs, all tagged genes were under the control of their endogenous promoter. Successful integration was confirmed by a combination of fluorescent microscopy, PCR and DNA sequencing.

Cells were grown to exponential phase for 36–48 hours before imaging. Strains containing the *Pnmt81-GFP-atb2* and *Pnmt41-GFP-CHD* constructs were shifted to EMM5S 15-18 hours prior to imaging to allow expression of the construct. To depolymerize the microtubules, 50 ug/mL MBC (methyl-2-benzimidazole-carbamate) was added to the cells immediately before imaging (Sawin and Snaith, 2004; Castagnetti *et al*., 2010).

### Spinning-Disk Confocal Microscopy

Cells were grown to exponential phase at 25 °C in YE5S-rich liquid medium in 50-mL baffled flasks in a shaking temperature-controlled incubator in the dark. Cells were concentrated 10- to 20-fold by centrifugation at 2,400 × g for 30 s and then resuspended in EMM5S. 5 μL of cells were mounted on a thin gelatin pad consisting of 10 μL 25% gelatin (Sigma Aldrich; G-2500) in EMM5S, sealed under a #1.5 coverslip with VALAP (1:1:1Vaseline:Lanolin:Parafin) and observed at 22 °C.

Fluorescence micrographs of live cells were acquired with a Nikon Eclipse Ti microscope equipped with a 100×/numerical aperture (NA) 1.49 HP Apo TIRF objective (Nikon), a CSU-X1 (Yokogawa) confocal spinning-disk system, 488/561 nm solid state lasers, and an electron-multiplying cooled charge-coupled device camera (EMCCD IXon 897, Andor Technology). The Nikon Element software was used for acquisition.

ImageJ (Schneider *et al*., 2012) was used for all image visualization and analyses. Images in the figures are maximum intensity projections of z-sections spaced at 0.36 μm. Images were systematically contrasted to provide the best visualization, and images within the same figure panel were contrasted using the same settings.

### Image analyses

The relative position of the nucleus (Figure 1B and Supplemental Figure 4A) was calculated by measuring the length of the cell compartment the nucleus inhabited (L_C_) and the distance of the center of the nucleus from the cell end (L_E_). The center of the nucleus was determined by eye using high magnification images. The radius of each nucleus (R_N_) was assumed to be roughly 1 μm (diameter, 2R_N_, of 2 μm) based on our measurements in wild-type cells. As the center of the nucleus cannot occupy either the very cell tip or the center of the contractile ring, we calculated the travelable distance as the diameter (2R_N_) of the nucleus subtracted from the cell compartment length. Using this travelable distance, these values were then normalized so that a value of 0.00 represented the closest a nucleus of could get to the end, while 1.00 represented the closest a nucleus could get to the contractile ring. This was calculated for each value with the following formula:

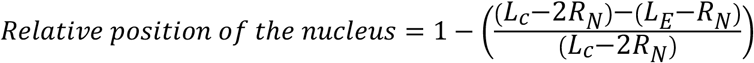

Cells expressing mEGFP-Myo2p Cut11p-tdTomato were used to calculate the relative position of the nucleus for all time points including and prior to 10 minutes after septation, while cells expressing GFP-CHD Cut11p-tdTomato were used to calculate the relative position of the nucleus for all time points occurring later than 10 minutes after septation.

Timing of microtubule events (Figure 2E) was measured by noting the camera frame when the spindle appeared (t=0), the spindle reached the cell ends, the spindle broke down, the ePAA appeared, the lPAA appeared, the pre-interphase microtubules appeared, and the mSPBs microtubules appeared. The spindle was considered fully elongated when the ends of the spindle no longer progressed towards the cell end. The spindle was considered broken down when it was no longer a continuous, single entity after the completion of spindle elongation.

The timing of events associated with the nucleus (Figure 2F) was measured by noting the first camera frame when the nucleus reached the cell end, the SPB faced the cell end, the completion of SPB rotation, and the completion of the displacement of the nucleus after spindle disassembly.

Contractile ring timing (Figure 2F) was measured by noting the camera frame when the spindle pole bodies separated (t=0) and the contractile ring disassembled. SPB separation was defined as the time when the spindle pole bodies were first distinguishable as two objects. Completion of contractile ring disassembly was defined as the time when the signal from the constricting contractile ring was no longer visible at the end of constriction.

To track the position of the nucleus (Figure 3C), the coordinates of the position of the end of the nucleus closest to the contractile ring were measured every minute from the time that the nucleus reached the cell end until the end of ring disassembly. From these measurements, the displacement velocity (Supplemental Figure 2) was calculated by taking the change in position at the start of displacement versus the end of displacement and dividing this value by the duration of displacement.

The position of the SPB over time during reorientation (Figure 4C) was tracked using the TrackMate plugin in FIJI (Tinevez *et al*., 2017). The coordinates of the traces were corrected for the vertical change in position of the nucleus at each time point to account for the displacement away from the cell end that the nucleus undergoes after spindle disassembly.

The extent of SPB rotation (Figure 4D) was calculated by determining whether the SPB rotated from facing the cell end at time of spindle disassembly to occupying the hemisphere of the nucleus facing the contractile ring before completion of septation.

The interactions of the lPAA microtubule +tips with the nucleus (Figure 5F and G) were measured by merging channels containing the Cut11p-tdTomato and the GFP-Atb2p (or Mal3p-mEGFP) markers. After spindle disassembly but before septation, the number of times GFP-Atb2p colocalized with the Cut11p-tdTomato marker was measured every 60 seconds. The duration of each colocalization event was also measured to determine how stable these interactions were.

To generate traces of the position of the proximal end of the mSPB microtubules over time (Figure 5H), the coordinates of the position of the proximal ends of the mSPB microtubules were determined every minute from the time of appearance to ring disassembly. We then calculated the percentage of contacts with the hemispherical cell end (within 1 μm of the cell end) and the lateral (> 1 μm away from the cell end) portion of the cell edge for the first and second half of all the contacts.

The elongation and shrinkage velocities of the lPAA microtubules were calculated from the change in position of the lPAA microtubules from the beginning to the end of elongation or shrinkage and dividing this value by the duration of either elongation or shrinkage.

### Computer Simulations

Simulations were run using the CytoSim platform (Nédélec and Foethke, 2007), a Brownian Dynamics simulator where each component of the cytoskeleton is represented as an individual object with associated physical and interactive properties and subjected to thermal fluctuations. The configuration file from Foethke et al., 2009 was modified to allow for asymmetric cell growth (see config.cym file) (Foethke *et al*., 2009). The 2009 platform, with hemispherical cell ends, does not allow for asymmetric cell growth. To overcome this challenge, we simulated the cell as a long cylinder with two large beads to either side of the cell to demarcate the cell ends. The beads were 100 μm in radius and had a viscosity of 10,000 pN*s/μm^2^, so that the force exerted by the microtubules did not push the cell ends. In simulations where growth was considered, the bead representing the old end was moved at a rate of 0.024 μm/min after 10 min of simulation time (OETO), and the new end moved in the opposite direction at a rate of 0.024 μm/min after 60 min (NETO) of simulation time, to reflect our experimental measurements in wild-type cells. At each timepoint in the simulation, we extracted the positions of the old end, the new end, the center of the nucleus, and the +ends of the interphase microtubules. The +tips of the microtubules were considered to contact the cell ends if they came within 0.1 μm of the beads. Significance in the difference of final nuclear position at 30-40 minutes after start of the simulation between the types of simulations was calculated using Tukey’s HSD. All parameters used in the configuration file are listed in Supplemental Table 2.

All simulations start with *de novo* microtubule polymerization from sites around the nuclear envelope. This initial growth resulted in the initial 80 s profile of nuclear displacement visible in all simulations. All nuclear displacement traces (Figure 7A-D) show the position of the center of the nucleus from the proximal cell end over time. The relative position of the nucleus was calculated as previously described for experimental data acquired in live cells (see above). To calculate the peak velocity, we measured the change in the position over time during the period of fastest movement at the beginning of displacement using the average of the displacement traces from all simulations within a dataset (Figure 6A). To calculate the average velocity, we measured the change in the position over time during the entire period of nuclear movement using the average of the displacement traces from all simulations within a dataset (Figure 6A). To calculate Δt_d-p,_ we measured the time required for either the proximal or distal end of the microtubules to reach either the proximal (Δt_p_) or distal end (Δt_d_) of the cell respectively; we then subtracted Δt_p_ from Δt_d_ to determine the additional time needed to reach the distal end (Figure 6B and C). To calculate Δc_p-d, we_ measured the total number of microtubules contacts at either the proximal (Δc_p_) or distal end (Δc_d_) of the cell during the period before the nucleus reached a stable position within the cell (first 20 min of simulation). We then subtracted Δc_d_ from Δc_p_ to determine the additional contacts present at the proximal end. We were interested in Δc_p-d_ only during the period of nuclear displacement as this is when presence of a net pushing force would be most meaningful.

## FIGURE LEGENDS

**Supplemental Figure 1.**
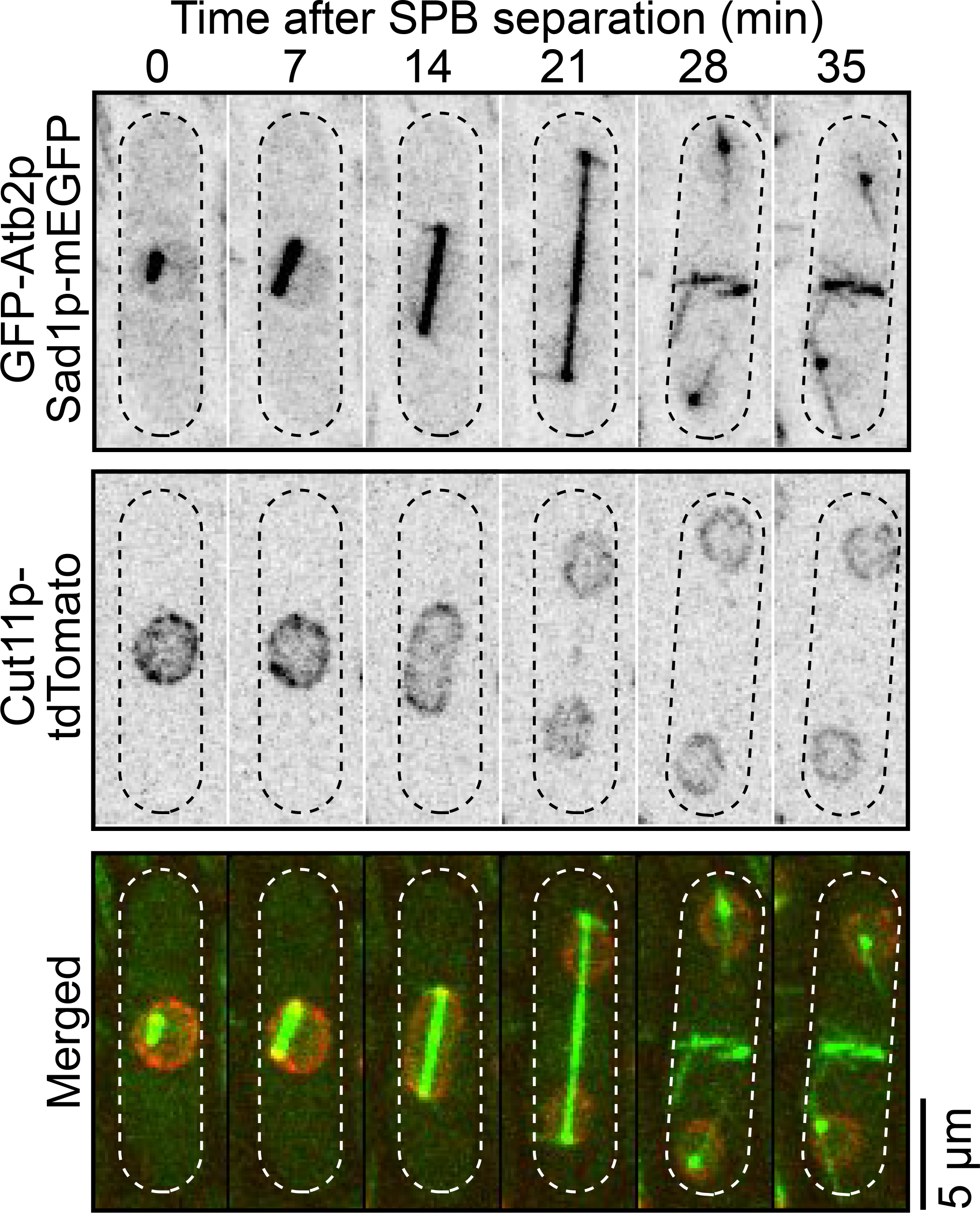
Position of daughter during anaphase. Time series micrographs of a wild-type cell showing daughter nuclei separation by the elongating spindle. Inverted grayscale for single color, merged for two colors and dotted are cell outlines.

**Supplemental Figure 2.**
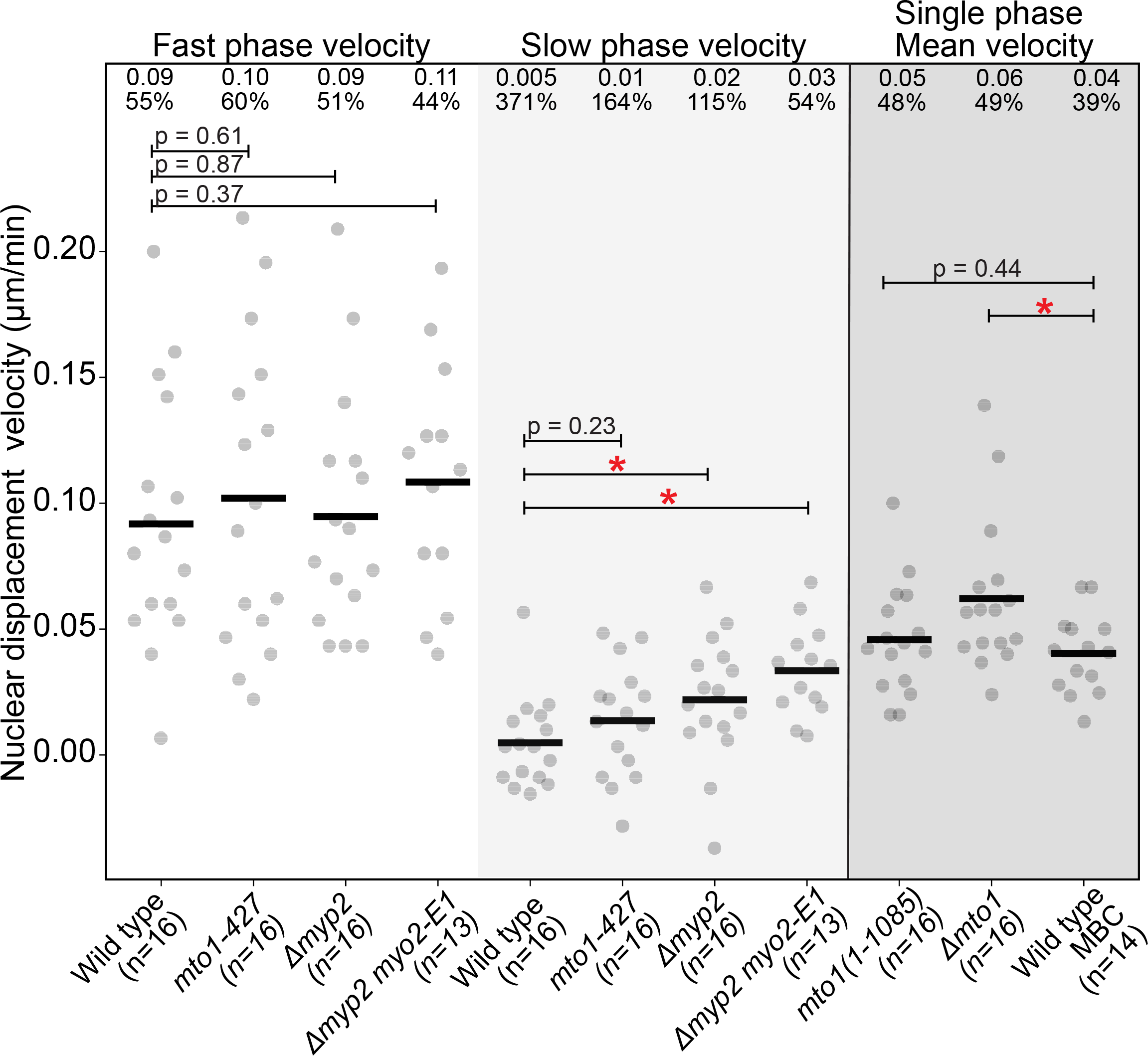
mSPB microtubules are essential for nuclear displacement before septation. Swarm plot of the nuclear displacement velocities from Figure 3C. For genotypes with PAA microtubules, velocities were calculated for both the fast and the slow displacement phases. For genotypes that lacked PAA microtubules, a single velocity for the entire displacement was calculated. Mean velocity and coefficient of variation are listed at the top of the plot for each dataset. Red asterisks, p < 0.05 by Student’s t-test. Bars, means.

**Supplemental Figure 3.**
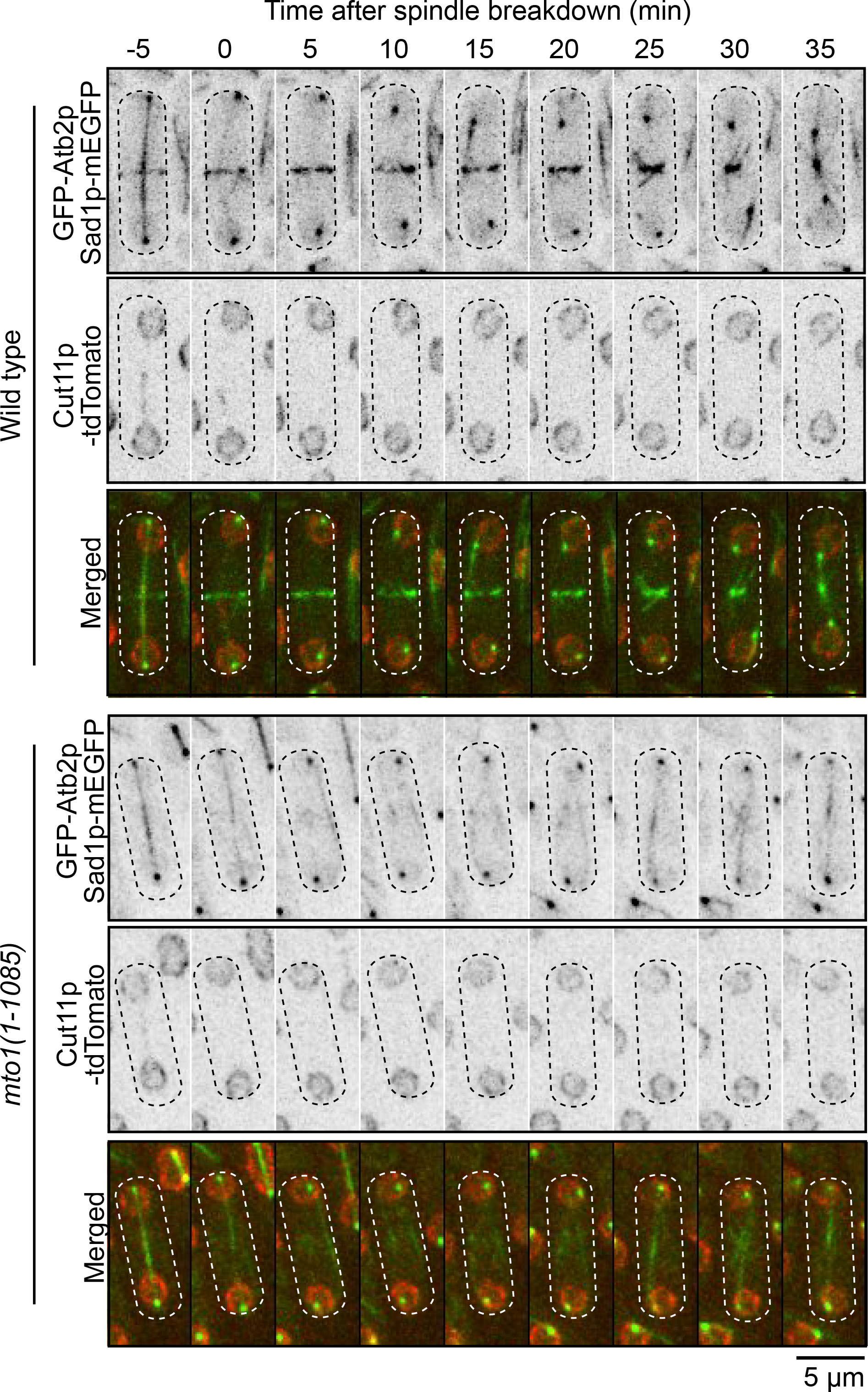
mSPB microtubules rotate the nucleus. Time series of micrographs of a wild-type and *mto1(1-1085)* cell showing the rotation (or lack of rotation) of the SPB. All micrographs, inverted grayscale for single color, merged for two colors and dotted lines are cell outlines.

**Supplemental Figure 4.**
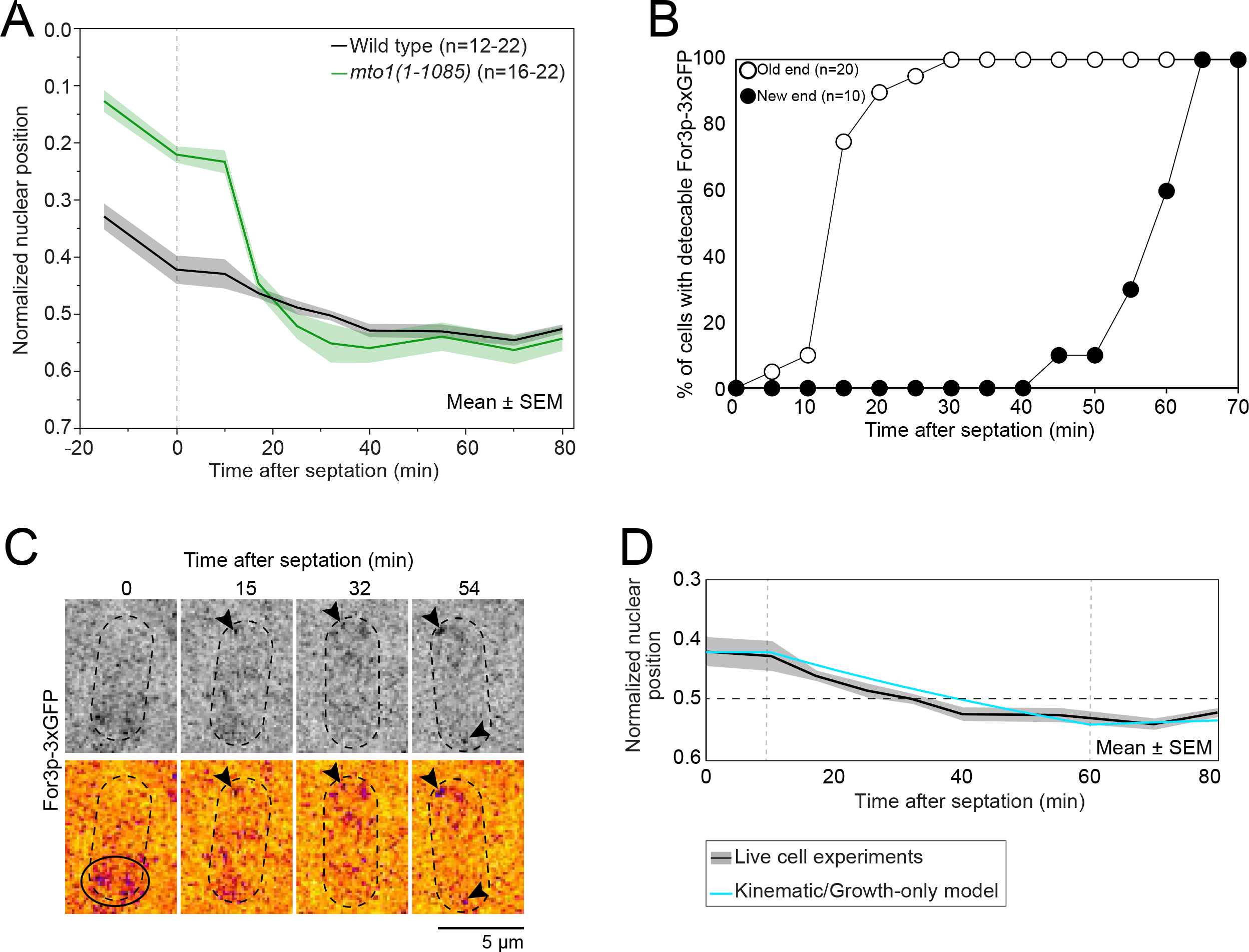
Asymmetric cell growth influences nuclear displacement. **A**. Plot of the nuclear position within the cell, normalized for the cell length. 0, closest distance to old cell end. 1, closest distance to new cell end. Vertical dashed line, time of septation **B**. Outcomes plot showing the timing of arrival of For3-3xGFP after septation to either the old end or new end of the cell. Time zero, septation. **C**. Time series micrographs showing the appearance of For3-3xGFP to the old end and the new end after septation. Inverted grayscale, top. Fire, bottom. Arrowhead, For3-3xGFP spots. Circle, For3-3xGFP signal present at new end at septation is disappearing remnants from disassembled ring. **D**. Plot of the relative position of the nucleus over time measured experimentally in live cells (same as in Supplemental Figure 4A) and calculated by the kinematic/growth-only model of a cell with an immobile nucleus. Vertical grey dashed lines, time of OETO and NETO respectively.

**Supplemental Video 1. Simulated nuclear displacement in 12.4-μm cell**. Video output of simulation of nuclear displacement in a 12.4 μm cell with a nucleus positioned at the cell end.

**Supplemental Video 2. Simulated nuclear displacement in growing 7-μm cell**. Video output of simulation of nuclear displacement in a 7 μm cell with a nucleus positioned at a P/D ratio of 0.83 to mimic a newborn cell. OETO starts at 10 min and NETO at 60 min.

## ACKNOWLEDGEMENTS

We thank Kenneth Sawin, Samantha Dundon, Thomas D. Pollard for strains and valuable input on this work. This research was supported by National Institute of Health Research Grant 5R01GM134254 and an internal grant from the college of veterinary medicine at NCSU. The authors declare no competing financial interests.

---

**Supplemental Table 1.**

| Strain | Genotype | Reference/Origin |
| --- | --- | --- |
| CL380 | *hphMX6-nmt81-GFP-atb2 kanMX6-Pmyp2sh-mCherry-myp2* | This work. |
| CL563 | *h+ kanMX6-Pmyo2short-mEGFP-myo2 sad1-RFP-kanMX6* | Bellingham-Johnstun et al., 2021 |
| CL635 | *h- for3-3GFP-ura4+ sad1-RFP-kanMX6* | Bellingham-Johnstun et al., 2021 |
| CL647 | *h- cut11-tdTomato-natMX6 hphMX6-nmt81-GFP-atb2 rlc1-tdTomato-natMX6 sad1-mEGFP-kanMX6* | This work. |
| CL676 | *h+ mal3-mEGFP-kanMX6 cut11-tdTomato-natMX6 rlc1-tdTomato-natMX6* | This work. |
| CL719 | *h- Δmto1::kanMX6 hphMX6-nmt81-GFP-atb2 cut11-tdTomato-natMX6 sad1-mEGFP-kanMX6* | This work. |
| CL720 | *Δmto1::kanMX6 kanMX6-Pmyo2short-mEGFP-myo2 cut11-tdTomato-natMX6 sad1-RFP-kanMX6* | This work. |
| CL721 | *h- kanMX6-Pmyo2short-mEGFP-myo2 cut11-tdTomato-natMX6 sad1-RFP-kanMX6* | This work. |
| CL751 | *cut11-tdTomato-natMX6 hphMX6-nmt81-GFP-atb2 sad1-mEGFP-kanMX6* | This work. |
| CL754 | *h+ mto1-427 cut11-tdTomato-natMX6 hphMX6-nmt81-GFP-atb2 sad1-mEGFP-kanMX6* | This work. |
| CL756 | *h- mto1-427 kanMX6-Pmyo2short-mEGFP-myo2 cut11-tdTomato-natMX6 sad1-RFP-kanMX6* | This work. |
| CL777 | *mto1(1-1085):ura4+ kanMX6-Pmyo2sh-mEGFP-myo2 cut11-tdTomato-natMX6 sad1-RFP-kanMX6* | This work. |
| CL778 | *mto1(1-1085):ura4+ cut11-tdTomato-natMX6 hphMX6-nmt81-GFP-atb2 sad1-mEGFP-kanMX6* | This work. |
| CL794 | *h+ Δmyp2::his7+ kanMX6-Pmyo2sh-mEGFP-myo2 cut11-tdTomato-natMX6 sad1-RFP-kanMX6* | This work. |
| CL795 | *h- Δmyp2::his7+ cut11-tdTomato-natMX6 hphMX6-nmt81-GFP-atb2 sad1-mEGFP-kanMX6* | This work. |
| CL823 | *Δmyp2::his7+ kanMX6-Pmyo2short-mEGFP-myo2-E1 cut11-tdTomato-natMX6* | This work. |
| CL824 | *Δmyp2::his7+ myo2-E1 cut11-tdTomato-natMX6 sad1-mEGFP-KanMX6* | This work. |
| CL848 | *nmt41-GFP-CHD-leu1+ rlc1-tdTomato-natMX6 cut11-tdTomato-natMX6* | This work. |
| CL850 | *mto1-427 nmt41-GFP-CHD-leu1+ cut11-tdTomato-natMX6* | This work. |
| CL852 | *mto1(1-1085):ura4+ nmt41-GFP-CHD-leu1+ rlc1-tdTomato-natMX6 cut11-tdTomato-natMX6* | This work. |

**Supplemental Table 2.**
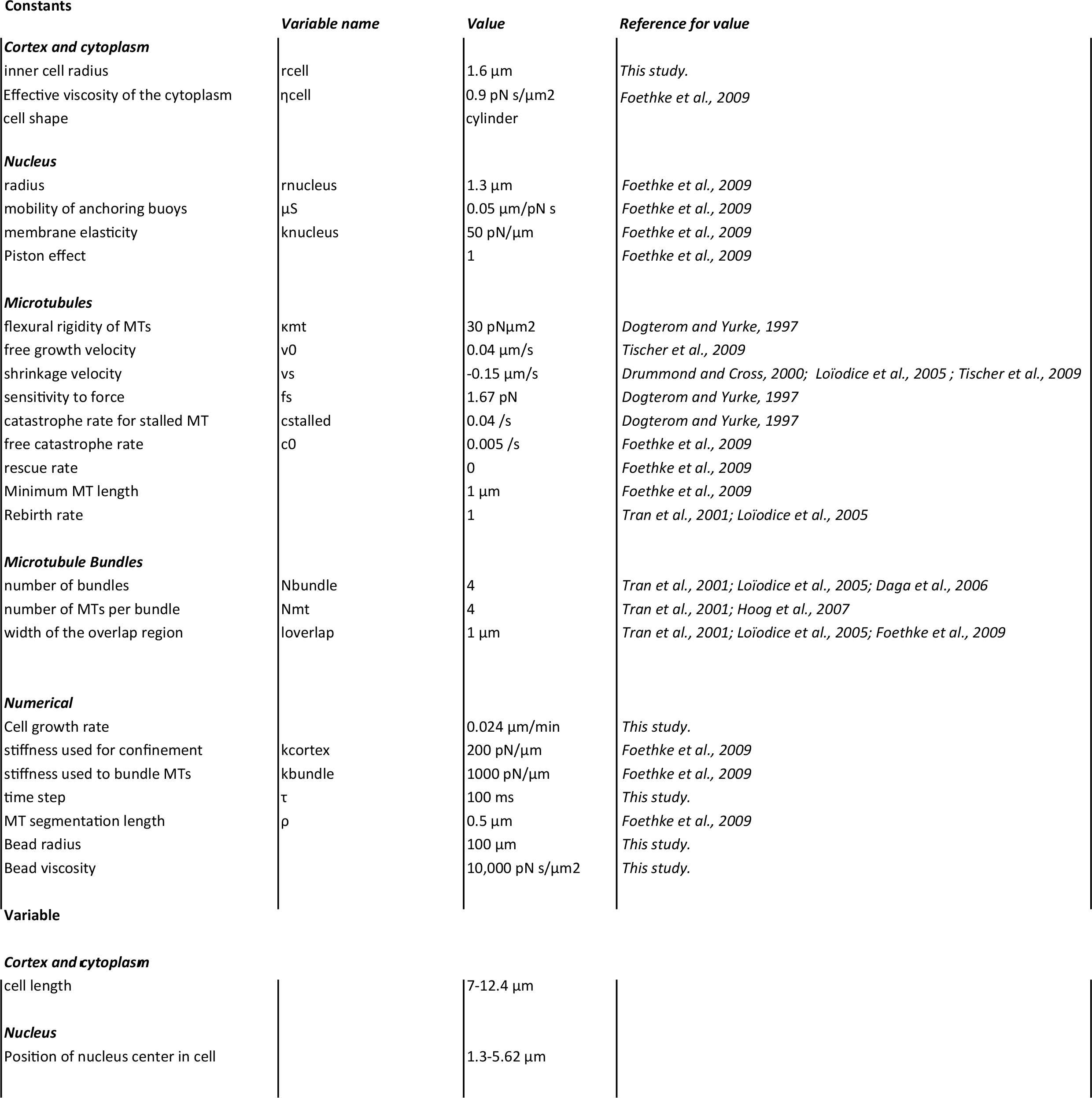

## REFERENCES

Akamatsu, M., Berro, J., Pu, K.M., Tebbs, I.R., and Pollard, T.D. (2014). Cytokinetic nodes in fission yeast arise from two distinct types of nodes that merge during interphase. J Cell Biol 204, 977–988.

Bahler, J., Wu, J.Q., Longtine, M.S., Shah, N.G., McKenzie, A., 3rd, Steever, A.B., Wach, A., Philippsen, P., and Pringle, J.R. (1998). Heterologous modules for efficient and versatile PCR-based gene targeting in Schizosaccharomyces pombe. Yeast 14, 943–951.

Browning, H., Hayles, J., Mata, J., Aveline, L., Nurse, P., and McIntosh, J.R. (2000). Tea2p is a kinesin-like protein required to generate polarized growth in fission yeast. J Cell Biol 151, 15–28.

Brunner, D., and Nurse, P. (2000). CLIP170-like tip1p spatially organizes microtubular dynamics in fission yeast. Cell 102, 695–704.

Castagnetti, S., Oliferenko, S., and Nurse, P. (2010). Fission yeast cells undergo nuclear division in the absence of spindle microtubules. PLoS Biol 8, e1000512.

Daga, R.R., and Chang, F. (2005). Dynamic positioning of the fission yeast cell division plane. Proc Natl Acad Sci U S A 102, 8228–8232.

Daga, R.R., Yonetani, A., and Chang, F. (2006). Asymmetric microtubule pushing forces in nuclear centering. Curr Biol 16, 1544–1550.

Ding, R., McDonald, K.L., and McIntosh, J.R. (1993). Three-dimensional reconstruction and analysis of mitotic spindles from the yeast, Schizosaccharomyces pombe. J Cell Biol 120, 141–151.

Ding, R., West, R.R., Morphew, D.M., Oakley, B.R., and McIntosh, J.R. (1997). The spindle pole body of Schizosaccharomyces pombe enters and leaves the nuclear envelope as the cell cycle proceeds. Mol Biol Cell 8, 1461–1479.

Feierbach, B., Verde, F., and Chang, F. (2004). Regulation of a formin complex by the microtubule plus end protein tea1p. J Cell Biol 165, 697–707.

Felgner, H., Frank, R., Biernat, J., Mandelkow, E.M., Mandelkow, E., Ludin, B., Matus, A., and Schliwa, M. (1997). Domains of neuronal microtubule-associated proteins and flexural rigidity of microtubules. J Cell Biol 138, 1067–1075.

Foethke, D., Makushok, T., Brunner, D., and Nedelec, F. (2009). Force- and length-dependent catastrophe activities explain interphase microtubule organization in fission yeast. Mol Syst Biol 5, 241.

Gundersen, G.G., and Worman, H.J. (2013). Nuclear positioning. Cell 152, 1376–1389.

Hagan, I., and Yanagida, M. (1997). Evidence for cell cycle-specific, spindle pole body-mediated, nuclear positioning in the fission yeast Schizosaccharomyces pombe. J Cell Sci 110 (Pt 16), 1851–1866.

Hagan, I.M., and Hyams, J.S. (1988). The use of cell division cycle mutants to investigate the control of microtubule distribution in the fission yeast Schizosaccharomyces pombe. J Cell Sci 89 (Pt 3), 343–357.

Heitz, M.J., Petersen, J., Valovin, S., and Hagan, I.M. (2001). MTOC formation during mitotic exit in fission yeast. J Cell Sci 114, 4521–4532.

Laplante, C., Berro, J., Karatekin, E., Hernandez-Leyva, A., Lee, R., and Pollard, T.D. (2015). Three myosins contribute uniquely to the assembly and constriction of the fission yeast cytokinetic contractile ring. Curr Biol 25, 1955–1965.

Loiodice, I., Staub, J., Setty, T.G., Nguyen, N.P., Paoletti, A., and Tran, P.T. (2005). Ase1p organizes antiparallel microtubule arrays during interphase and mitosis in fission yeast. Mol Biol Cell 16, 1756–1768.

Mana-Capelli, S., McLean, J.R., Chen, C.T., Gould, K.L., and McCollum, D. (2012). The kinesin-14 Klp2 is negatively regulated by the SIN for proper spindle elongation and telophase nuclear positioning. Mol Biol Cell 23, 4592–4600.

Martin, S.G., and Chang, F. (2005). New end take off: regulating cell polarity during the fission yeast cell cycle. Cell Cycle 4, 1046–1049.

Martin, S.G., McDonald, W.H., Yates, J.R., and Chang, F. (2005). Tea4p links microtubule plus ends with the formin for3p in the establishment of cell polarity. Dev Cell 8, 479–491.

McCully, E.K., and Robinow, C.F. (1971). Mitosis in the fission yeast Schizosaccharomyces pombe: a comparative study with light and electron microscopy. J Cell Sci 9, 475–507.

Mickey, B., and Howard, J. (1995). Rigidity of microtubules is increased by stabilizing agents. J Cell Biol 130, 909–917.

Mitchison, J.M., and Nurse, P. (1985). Growth in cell length in the fission yeast Schizosaccharomyces pombe. J Cell Sci 75, 357–376.

Nédélec, F., and Foethke, D. (2007). Collective Langevin dynamics of flexible cytoskeletal fibers. New Journal of Physics 9.

Pardo, M., and Nurse, P. (2003). Equatorial retention of the contractile actin ring by microtubules during cytokinesis. Science 300, 1569–1574.

Pichova, A., Kohlwein, S.D., and Yamamoto, M. (1995). New Arrays of Cytoplasmic Microtubules in the Fission Yeast Schizosaccharomyces-Pombe. Protoplasma 188, 252–257.

Samejima, I., Miller, V.J., Rincon, S.A., and Sawin, K.E. (2010). Fission yeast Mto1 regulates diversity of cytoplasmic microtubule organizing centers. Curr Biol 20, 1959–1965.

Sawin, K.E., and Snaith, H.A. (2004). Role of microtubules and tea1p in establishment and maintenance of fission yeast cell polarity. J Cell Sci 117, 689–700.

Schneider, C.A., Rasband, W.S., and Eliceiri, K.W. (2012). NIH Image to ImageJ: 25 years of image analysis. Nat Methods 9, 671–675.

Snaith, H.A., Samejima, I., and Sawin, K.E. (2005). Multistep and multimode cortical anchoring of tea1p at cell tips in fission yeast. EMBO J 24, 3690–3699.

Snaith, H.A., and Sawin, K.E. (2003). Fission yeast mod5p regulates polarized growth through anchoring of tea1p at cell tips. Nature 423, 647–651.

Tanaka, K., and Kanbe, T. (1986). Mitosis in the fission yeast Schizosaccharomyces pombe as revealed by freeze-substitution electron microscopy. J Cell Sci 80, 253–268.

Tinevez, J.Y., Perry, N., Schindelin, J., Hoopes, G.M., Reynolds, G.D., Laplantine, E., Bednarek, S.Y., Shorte, S.L., and Eliceiri, K.W. (2017). TrackMate: An open and extensible platform for single-particle tracking. Methods 115, 80–90.

Tolic-Norrelykke, I.M., Sacconi, L., Stringari, C., Raabe, I., and Pavone, F.S. (2005). Nuclear and division-plane positioning revealed by optical micromanipulation. Curr Biol 15, 1212–1216.

Tran, P.T., Doye, V., Chang, F., and Inoue, S. (2000). Microtubule-dependent nuclear positioning and nuclear-dependent septum positioning in the fission yeast Schizosaccharomyces [correction of Saccharomyces] pombe. Biol Bull 199, 205–206.

Tran, P.T., Marsh, L., Doye, V., Inoue, S., and Chang, F. (2001). A mechanism for nuclear positioning in fission yeast based on microtubule pushing. J Cell Biol 153, 397–411.

Venkatram, S., Tasto, J.J., Feoktistova, A., Jennings, J.L., Link, A.J., and Gould, K.L. (2004). Identification and characterization of two novel proteins affecting fission yeast gamma-tubulin complex function. Mol Biol Cell 15, 2287–2301.

Verde, F., Mata, J., and Nurse, P. (1995). Fission yeast cell morphogenesis: identification of new genes and analysis of their role during the cell cycle. J Cell Biol 131, 1529–1538.

Yukawa, M., Teratani, Y., and Toda, T. (2021). Escape from mitotic catastrophe by actin-dependent nuclear displacement in fission yeast. iScience 24, 102031.

Zhang, D., Vjestica, A., and Oliferenko, S. (2010). The cortical ER network limits the permissive zone for actomyosin ring assembly. Curr Biol 20, 1029–1034.

Zimmerman, S., and Chang, F. (2005). Effects of {gamma}-tubulin complex proteins on microtubule nucleation and catastrophe in fission yeast. Mol Biol Cell 16, 2719–2733.

